# Pupil-linked arousal differentially modulates cell-type-specific sensory processing

**DOI:** 10.1101/2025.06.09.658645

**Authors:** Keith J. Kaufman, Rebecca F. Krall, Ross S. Williamson

## Abstract

Arousal is a ubiquitous influence on the brain that fluctuates during wakefulness and modulates sensation and perception. These fluctuations impact membrane potential, cortical state, and sensory encoding, yet prior studies report inconsistent effects, likely due to averaging across heterogeneous cell types. To resolve this, we combined two-photon calcium imaging and pupillometry in awake mice to examine arousal-related activity in excitatory subpopulations of the auditory cortex: intratelencephalic (IT), extratelencephalic (ET), and corticothalamic (CT) neurons. Pupil-linked arousal modulated all cell types through diverse linear and non-linear response motifs. ET neurons showed significant multiplicative and additive gain modulation, with enhanced response magnitude and encoding but reduced frequency selectivity. CT and L2/3 neurons exhibited inverted-U relationships between arousal and both response strength and decoding accuracy, while IT neurons were minimally affected. These patterns closely tracked changes in population-level reliability, revealing a mechanistic link between internal state and the stability of sensory representations.

## Introduction

The neural basis of perception involves the interplay between external stimuli and internal states. Internal state refers to a range of physiological and psychological variables, including hunger, emotion, and arousal, that dynamically shape sensory processing [1–3]. Changes in brain state, from deep sleep to heightened arousal, can profoundly influence how sensory information is encoded and processed [4–6]. Yet, while state transitions reshape neural activity across the brain, their specific effects on distinct cortical cell types remain poorly understood.

Autonomic fluctuations in pupil diameter offer a non-invasive, real-time index of internal state fluctuations [7, 8]. Far from serving solely as an aperture for light, the pupil reflects a wide array of non-luminance-linked processes, including attention, cognitive effort, imagined illumination, arousal, and even heart rate [7, 9–19]. Pupil diameter fluctuations co-vary with activity in several neuromodulatory systems that regulate waking states, such as the locus coeruleus (norepinephrine), basal forebrain (acetylcholine), and dorsal raphe (serotonin) systems [12, 20–33]. These systems project broadly to cortex, where they modulate excitability, network dynamics, and cognition, supporting the view that pupil size serves as a proxy for neuromodulatory tone.

Consistent with this view, pupil-linked arousal has been shown to modulate cortical membrane potentials, spontaneous and evoked firing, tuning selectivity, stimulus encoding, and pairwise neural correlations across sensory areas [9, 11, 34–36]. However, these effects are variable across studies, with both linear and non-linear relationships reported between arousal and neural activity [9, 11, 35, 37–39]. In visual (VCtx) and auditory (ACtx) cortices, arousal typically enhances spontaneous and evoked responses [11, 35, 37, 39], yet peak activity has also been observed at intermediate pupil sizes, consistent with an inverted-U relationship that mirrors behavioral performance in sensoryguided tasks [9, 38]. While arousal sharpens orientation tuning in VCtx, its effect on frequency tuning in ACtx are less consistent, with reports of decreased or unchanged selectivity [11, 35, 36]. The impact of pupil state on stimulus decoding accuracy in ACtx has also been contested. One study found that decoding improved with arousal despite reduced tuning selectivity, attributed to a concurrent reduction in noise correlations [35]. In contrast, more recent work reported an inverted-U relationship between decoding accuracy and pupil diameter, driven in part by a reduction in neural variability [40]. Indeed, decoding performance has been shown to correlate strongly with population-level response reliability of stimulus-evoked responses [41].

Discrepancies across prior studies may stem from methodological differences but are likely compounded by the absence of cell-type-specific approaches capable of resolving distinct arousal effects within heterogeneous neural populations. For example, vasoactive intestinal peptide (VIP)- and somatostatin (SOM)-expressing interneurons exhibit opposing responses during low and high pupil-linked arousal states [11, 42]. Stimulus encoding recruits an array of distinct excitatory cell types that span the cortical lamina, including intratelencephalic (IT), extratelencephalic (ET), and corticothalamic (CT) cells (**Figure 1A**) [43, 44]. These subtypes differ in genetic profile, morphology, intrinsic excitability, and synaptic connectivity, all of which may shape their susceptibility to arousal-dependent modulation. IT cells are found in layers (L) 2-6 and project exclusively within the telencephalon, including the striatum, and ipsi-/ contralateral cortex [45, 46]. ET cells, located primarily in L5b, project to both telencephalic and subcortical targets, including the midbrain, thalamus, striatum, and amygdala [47, 48]. CT cells reside in L6, project locally to L5a, and send feedback connections to the thalamus [49–53]. Given this diversity, it is unlikely that arousal exerts uniform effects across excitatory subtypes. Instead, such heterogeneity may underlie the inconsistencies reported in previous studies.

**Figure 1:**
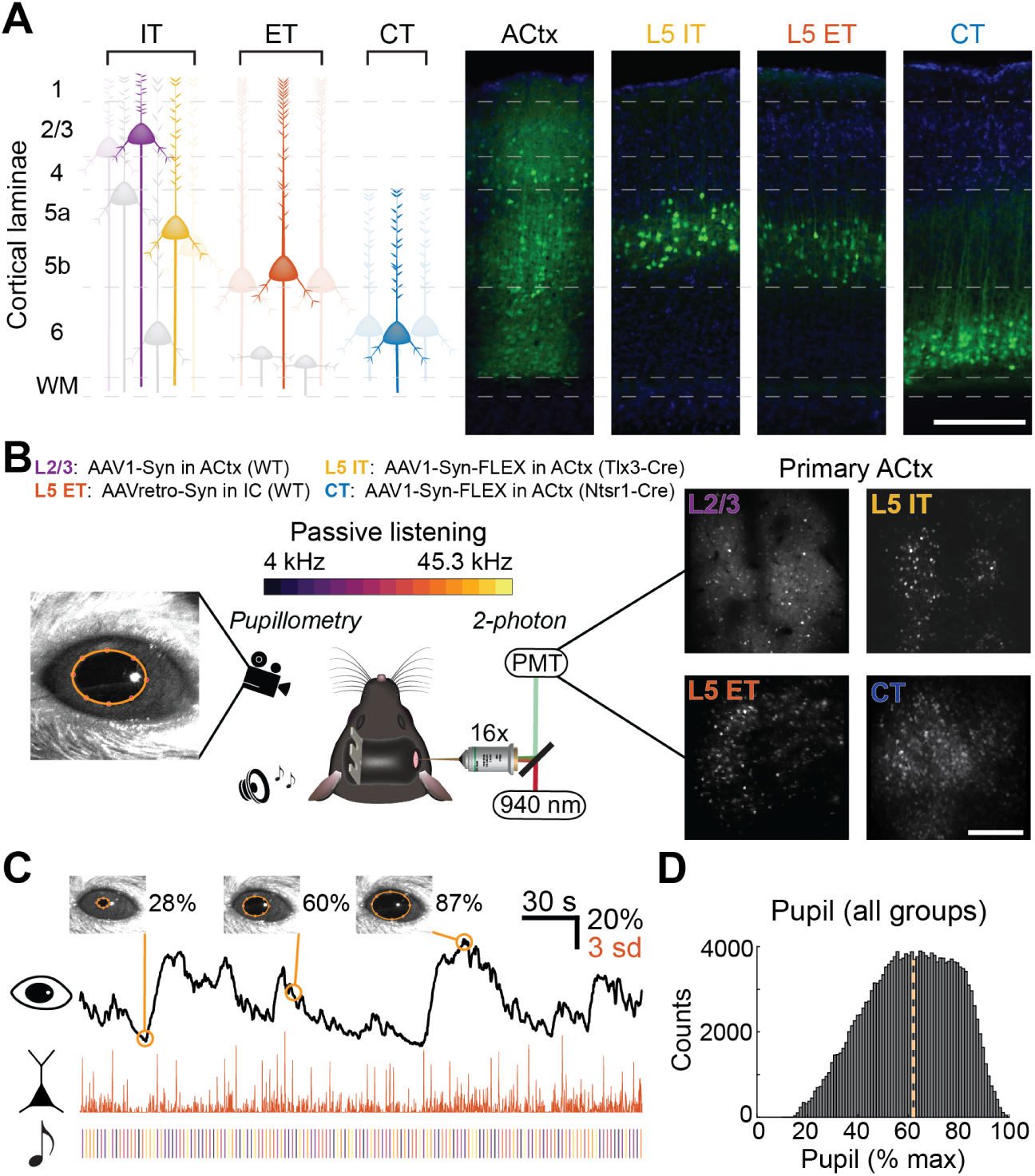
Experimental strategy for tracking pupil-linked arousal and cell-type-specific ACtx activity. **A**: Schematic of excitatory cortical projection neuron subtypes (left). Representative coronal sections of ACtx showing GCaMP8s expression (green) and DAPI-labeled (blue) nuclei (right). The transgenic and viral strategies noted in (B) were used to target L2/3, L5 intratelencenphalic (IT), L5 extratelencenphalic (ET), or corticothalamic (CT) neurons. Scale bar: 250 *µ*m. **B**: Overview of the experimental design. Mice passively listen to pure tone frequencies while two-photon calcium imaging and pupillometry were performed simultaneously. Cell-type-specific targeting strategies (top) used a combination of viral and/or transgenic methods (WT = wild type). Exemplar twophoton fields of view (FoV) of each target ACtx cell subpopulation (right). Scale bar: 250 *µ*m. **C**: Exemplar 5-minute recording showing pupil diameter (top, black), deconvolved neural activity from a single ET neuron (middle, orange), and sound onset times (bottom). Pupil size is expressed as a percentage of maximum diameter; neural activity is plotted in standard deviations (z-scored) from baseline. **D**: Histogram of trial-wise pupil states across all mice. Counts refer to number of trials. Median pupil size of 62% is denoted by the yellow dashed line.

To address this, we performed two-photon calcium imaging in awake mice to monitor the activity of IT, ET, and CT neurons in ACtx, while simultaneously tracking pupil diameter as a proxy for arousal. Each cell type exhibited a variety of pupil-associated response motifs, comprising both linear and non-linear shifts in activity. Arousal influenced frequency tuning selectivity across all subtypes, but L5 ET neurons uniquely showed multiplicative and additive transformations at heightened arousal levels. Although noise correlations decreased with pupil size across all groups, changes in signal correlations were cell-type-specific. Decoding analyses further revealed that pupil-linked arousal modulated stimulus classification accuracy in a cell-type-dependent manner, closely tracking changes in population-level reliability. Together, these findings demonstrate that arousal significantly shapes sensory encoding in ACtx via distinct mechanisms across excitatory subpopulations, providing a unifying framework to reconcile prior inconsistencies in the field.

## Results

To examine how arousal state influences sensory responses across excitatory subpopulations in ACtx, we simultaneously recorded neural activity and pupil diameter in awake, head-fixed mice. Animals passively listened to 15 pure tone frequencies (4 to ≈45.3 kHz at 70 dB SPL, 50 ms duration) presented in a pseudo-random order. We used two-photon calcium imaging to monitor neural activity in genetically defined excitatory cell-types in the right primary ACtx: L2/3 (*n* = 3, 023 neurons; *N* = 6 mice), L5 IT (*n* = 2, 447; *N* = 7), L5 ET (*n* = 2, 876; *N* = 10), and CT (*n* = 3, 311; *N* = 9). Each mouse expressed the genetically encoded calcium indicator, GCaMP8s, in a single subpopulation using a combination of viral and/or transgenic approaches (**Figure 1B**). L2/3 neurons were labeled via the injection of an AAV with a constitutive promoter into the ACtx of C57BL/6 mice. Cell-type-specific labeling of L5 IT and CT cells was accomplished via injection of a Cre-dependent viral vector in Tlx3 PL56-Cre and Ntsr1-Cre mice, respectively. L5 ET neurons were targeted via retrograde labeling from the inferior colliculus, a major projection target of ACtx ET neurons. Raw calcium signals were deconvolved and normalized to a 500 ms baseline window preceding sound onset to estimate relative spiking activity [54, 55]. In parallel, we tracked pupil diameter as a proxy for arousal (**Figure 1C**). Pupil size was normalized within each animal to its maximum observed diameter (across all imaging sessions). Pupil states on a given trial were computed by averaging the pupil trace during a baseline window (500 ms prior to sound onset). Baseline pupil diameters were used to characterize the instantaneous arousal state to account for slower pupil responses relative to neural activity. As in previous reports, pupil size distributions were approximately Gaussian, with most trials centered around an intermediate arousal level (median = 62%; **Figure 1D**). We therefore grouped pupil states into bins both above and below the median pupil diameter for subsequent analyses.

### Multivariate regression reveals diverse arousal and stimulus modulation motifs across excitatory subpopulations

We used a multivariate linear regression model to quantify how each neuron’s activity was modulated by sound and/or pupil-linked arousal. For each trial, a neuron’s evoked response was modeled as a function of stimulus identity and baseline neural, pupil, and pupil² activities (**Figure 2A**). This approach allowed us to estimate the proportion of response variance attributable to both sensory input and arousal, while capturing potential non-linear relationships via the quadratic pupil term. Regression coefficients for pupil and pupil² (*β_p_*, 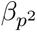) quantified the influence of linear and non-linear effects of arousal, respectively, while stimulus-related coefficients (*β_s_*) reflected tuning to individual frequencies. The *β_s_* values closely mirrored each neuron’s empirically measured tuning curve (**Figure 2B**). To determine the strength of this relationship, we computed the Pearson correlation between each neuron’s tuning curve and its *β_s_* weights, and compared the resulting distribution to a null distribution generated by shuffling stimulus labels for each cell. Observed correlations were significantly greater than chance (two-sample Kolmogorov–Smirnov test, *p <* 10^−100^), with the median Pearson’s *r* exceeding 0.99 across all cell types (**Supplementary Figure 1A**).

**Figure 2:**
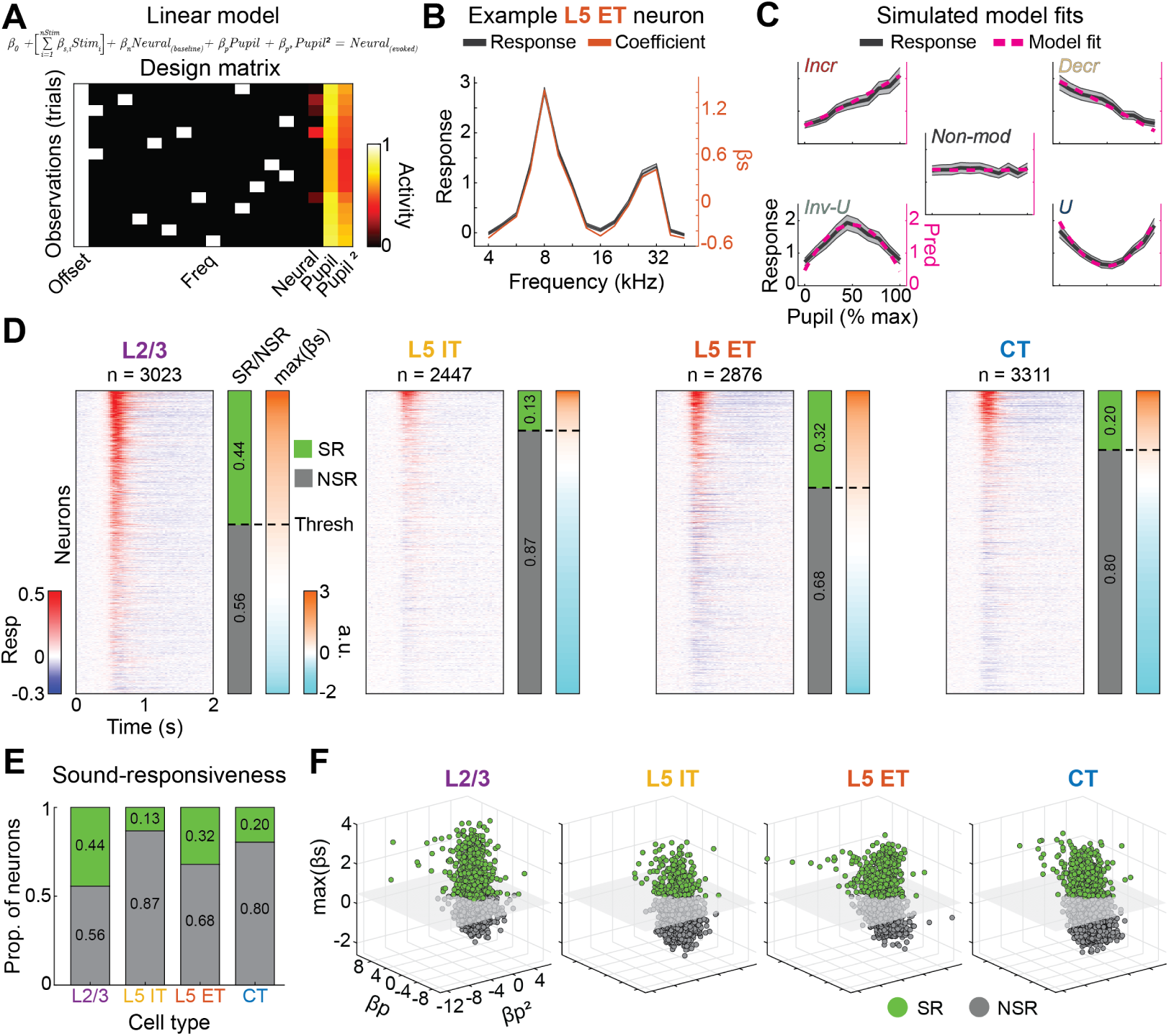
Multivariate regression reveals diverse arousal and stimulus modulation motifs across excitatory subpopulations. **A**: Truncated design matrix from a single L5 ET neuron used in the multivariate linear regression model (top). Predictors include one-hot encoded stimulus identity, and pre-stimulus baseline activity (neural, pupil, and pupil^2^). Normalized pupil activity is represented as a proportion of max dilation (range: 0–1). **B**: Frequency tuning curve (black) of the example L5 ET neuron in (A), overlaid with its corresponding stimulus-related regression coefficients (*β_s_*, orange). Shaded regions denote mean ± s.e.m. **C**: Simulated examples demonstrating five distinct arousal modulation motifs, each with identical frequency tuning but differing pupildependent responses. The full regression model accurately recovers each motif, including linear (increasing, decreasing), non-linear (inverted-U, U-shaped), and non-modulated profiles. Shaded regions denote mean ± s.e.m. **D**: Peristimulus time histograms (PSTHs) from all recorded neurons (collapsed across pupil state and stimulus), sorted within each subtype by their z-scored maximum *β_s_* value. Green indicates sound-responsive cells; gray indicates non-sound-responsive cells, based on a threshold of 0.5 standard deviations (dashed line). Cell counts: L2/3: *n* = 3, 023 neurons, *N* = 6 mice; L5 IT: *n* = 2, 447, *N* = 7; L5 ET: *n* = 2, 876, *N* = 10; CT: *n* = 3, 311, *N* = 9). **E**: Stacked bar plot showing the proportion of sound-responsive (green) and non-sound-responsive (gray) neurons across each excitatory subtype. Sound-responsiveness differs across excitatory subtype (Chi-square test, *χ*^2^ = 815.4, *p <* 10^−100^). **F**: 3D scatterplot of z-scored regression coefficients: *β_p_*, 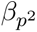, and maximum *β_s_*, for all neurons. Sound-responsive neurons (green) and non-sound-responsive neurons (gray) are separated by a threshold hyperplane along the *β_s_* axis.

To further validate our regression model, we simulated five neurons with identical frequency tuning but differing responses across pupil states; i.e., distinct state tuning curves (**Supplementary Figure 1B-D**). These synthetic neurons were designed to exhibit one of five pupil-dependent response patterns: increasing, decreasing, inverted-U (negative quadratic), U-shaped (positive quadratic), and non-modulated (**Figure 2C**). The full model, which included both linear and quadratic pupil terms, successfully captured each of these state tuning profiles. These results demonstrate that the model can accurately detect both linear and non-linear arousal modulation motifs.

Calcium imaging avoids the sampling bias inherent to electrophysiology, which tends to over-represent active neurons, making two-photon recordings well-suited for estimating the true proportion of sensory-responsive cells. To leverage this advantage, we used the regression model to assess each neuron’s sound responsiveness. For each neuron, we extracted the maximum stimulus-related *β_s_* coefficient (corresponding to its best frequency), and z-scored this value. Neurons with z-scored *β_s_* values exceeding 0.5 standard deviations were heuristically classified as sound-responsive, while all others were considered non-sound-responsive. This approach revealed significant differences in sound responsiveness across excitatory subtypes (**Figure 2D,E**, chi-squared test, *p <* 10^−100^). The highest proportion of sound-responsive neurons was observed in L2/3 (44%), followed by L5 ET (32%), CT (20%), and L5 IT (13%). On average, these sound-responsive neurons exhibited strong evoked activity across trials, irrespective of pupil state and stimulus (**Figure 2D**).

To visualize the diversity of pupil-related modulation across the entire dataset, we normalized each neuron’s *β_p_* and 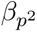 coefficients and plotted these alongside their maximum *β_s_* value (**Figure 2F**). If pupil state had no influence on sound-evoked activity, *β_p_* and 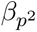 coefficients would cluster around zero. However, we observed a wide range of positive and negative coefficients, indicating that evoked responses were frequently modulated, both linearly and non-linearly, by pupil state. Together, these results demonstrate that pupil-linked arousal exerts a broad and heterogeneous influence on evoked neural activity across all layers and excitatory cell types of ACtx.

### Excitatory subtypes exhibit distinct patterns of arousal-dependent response modulation

The relationship between neural response magnitude and pupil-linked arousal has been debated in the literature [9, 11, 35, 37–39]. However, prior studies have not resolved how these relationships vary across excitatory neuron classes within a common cortical area. To address this, we compared arousal-dependent modulation of evoked responses across multiple excitatory subtypes in ACtx under identical stimulus conditions. For each neuron, we identified its best frequency (BF), defined as the pure tone frequency that elicited the greatest activity irrespective of pupil state, and quantified how BF-evoked activity varied with pupil diameter. These values were averaged within each pupil state bin to construct mean pupil-state tuning curves for each excitatory subtype (**Figure 3A**). Only sound-responsive neurons with sufficient data at enough pupil states were included (see Methods). Missing values for underrepresented bins were linearly interpolated or extrapolated according to defined inclusion criteria (**Supplementary Figure 2A-B**). Pupil-linked arousal significantly modulated response magnitude, and the pattern of modulation differed by excitatory subtype (**Figure 3A**, two-way mixed-effects model, main effect for pupil: *p* = 1.28 × 10^−10^, and cell type: *p <* 10^−100^, pupil x cell type interaction, *p* = 0.024). L2/3 (**Figure 3A**, FoVs = 21) and CT (**Figure 3A**, FoVs = 14) neurons exhibited inverted-U tuning curves, with maximal activity at intermediate arousal levels. In contrast, L5 ET neurons (**Figure 3A**, FoVs = 27) showed a linear increase in response magnitude with increasing pupil size. L5 IT neurons (**Figure 3A**, FoVs = 14) showed no modulation, suggesting that this subtype is relatively insensitive to arousal-linked changes in brain state.

**Figure 3:**
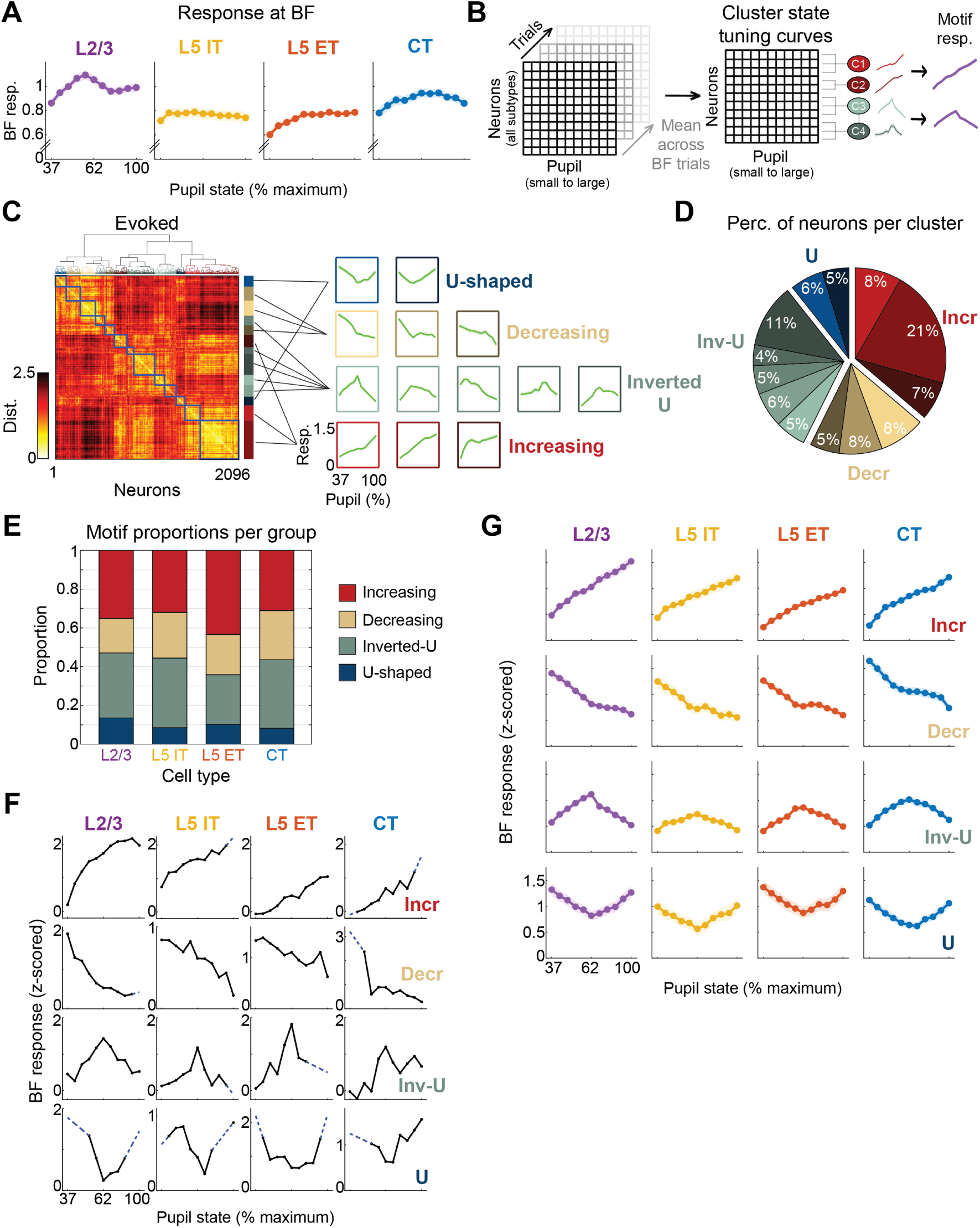
Excitatory subtypes exhibit distinct patterns of arousal-dependent response modulation. **A**: Mean best-frequency (BF) state tuning curves for each excitatory cell type; L2/3, *n* = 21 FoVs; L5 IT, *n* = 14 FoVs; L5 ET, *n* = 27 FoVs; CT, *n* = 14 FoVs; shaded regions, mean ± s.e.m.; two-way mixed-effects model, main effect for pupil and cell type, *F* = 6.77 and 235.73, *p* = 1.28 × 10^−10^ and *p <* 10^−100^, respectively; pupil x cell type interaction term, *F* = 1.57, *p* = 0.024. **B**: Schematic of unsupervised hierarchical clustering analysis of BF state tuning curves and manual grouping into functional pupil-state modulation motifs. **C**: Distance matrix from hierarchical clustering analysis of evoked state tuning curves (left). Mean response profiles for each cluster, organized by pupil-state modulation motifs (right). **D**: Pie chart showing the distribution of modulation motifs across clusters for all excitatory subpopulations. Colors correspond to cluster groupings in (C, right). Modulation motifs were not uniformly represented (*n* = 2096, chi-square goodness-of-fit test, *χ*^2^ = 330.3, *p* = 2.75 × 10^−71^). **E**: Proportions of pupil-state modulation motifs by excitatory cell-type. Motif distribution differed across groups (chi-square test of independence (*χ*^2^ = 43.59, *p* = 1.68 × 10^−6^). **F**: Representative BF state tuning curves from individual neurons for each motif and cell-type group. Dashed blue lines indicate linearly interpolated and/or extrapolated pupil-state data (see Methods). **G**: Mean BF state tuning curves by modulation motif and cell-type. Responses significantly differed across pupil state and cell type for increasing (*n* = 759, two-way ANOVA, main effect for pupil and cell type, *F* = 400 and 9.249, *p <* 10^−100^ and *p* = 5.18 × 10^−6^, respectively; cell type × pupil interaction term, *F* = 5.843, *p* = 3.77 × 10^−22^), decreasing (*n* = 443, two-way ANOVA, main effect for pupil and cell type, *F* = 268.7 and 3.418, *p <* 10^−100^ and *p* = 0.017, respectively; cell type × pupil interaction term, *F* = 2.876, *p* = 2.88 × 10^−7^), inverted-U (*n* = 670, two-way ANOVA, main effect for pupil and cell type, *F* = 122.8 and 3.472, *p <* 10^−100^ and *p* = 0.0159, respectively; cell type × pupil interaction term, *F* = 5.17, *p* = 1.51 × 10^−18^), and U-shaped motifs (*n* = 224, two-way ANOVA, main effect for pupil and cell type, *F* = 72.21 and 1.184, *p* = 9.16 × 10^−54^ and *p* = 0.317, respectively; cell type × pupil interaction term, *F* = 0.6606, *p* = 0.92). Shaded regions denote mean ± s.e.m.

Averaging the neural activity across large populations can obscure the diversity of statedependent response profiles. For instance, symmetric but opposing relationships (e.g., positive and negative quadratics) may cancel out when aggregated, resulting in a misleadingly flat average. To uncover hidden structure within the data, we applied hierarchical clustering to all sound-responsive neurons (**Figure 3B**). Clustering was performed separately using the neural activity from two distinct time windows: a baseline period (pre-stimulus onset) and an evoked period (post-stimulus onset). Dissimilarity between state tuning curves was quantified using Euclidean distance, and resulting clusters were classified into one of several functional pupil-state modulation motifs: increasing, decreasing, inverted-U (negative quadratic), or U-shaped (positive quadratic) (**Figure 3C**, **Supplementary Figure 2C-D**). While non-modulated neurons were present in both clustering analyses, pupil-modulated motifs were much more prominent. To ensure that any non-modulated cells scattered across the clusters did not distort the identified motifs, we confirmed that the mean z-scored tuning curve of each cluster closely matched the rescaled profile of its constituent cells (**Supplementary Figure 2E**). The distribution of all neurons across modulation motifs was significantly non-uniform in both the evoked (**Figure 3D**, chi-squared test, *n* = 2096, *p* = 2.75×10^−71^) and baseline (**Supplementary Figure 2D**, chi-squared test, *n* = 2096, *p <* 10^−100^) windows. While the four pupil-modulated motifs were found using both evoked and baseline activity, increasing (36%) and inverted-U (31%) motifs were most prevalent across evoked clusters, whereas increasing (49%) and decreasing (26%) motifs were most prevalent across baseline clusters (**Figure 3D** and **Supplementary Figure 2D**).

After assigning each neuron to a modulation motif via clustering, we examined how the distribution of these motifs varied across excitatory subtypes. Four pupil-state modulation motifs were represented within each subtype, but occurred at significantly different proportions (**Figure 3E**, chi-squared test, *p* = 1.68 × 10^−6^, **Supplementary Figure 2F-G**, chi-squared test, *p* = 0.015). The majority of sound-responsive neurons within each excitatory subtype exhibited either increasing responses or followed an inverted-U pattern with pupil dilation, consistent with previous findings [36]. The increasing motif was most prevalent in L2/3 and L5 ET neurons (35% and 43%, respectively), while the inverted-U motif was more common in L5 IT and CT populations (36% and 35%, respectively). The U-shaped motif was the least common across all cell types, observed in only 14% of L2/3, 8% of L5 IT, 10% of L5 ET, and 8% of CT cells. Overall, linear motifs (increasing or decreasing) were dominant over non-linear motifs (inverted-U or U-shaped) across all populations: L2/3 (53% linear vs. 47% non-linear), L5 IT (56% vs. 44%), L5 ET (64% vs. 36%), and CT (56% vs. 44%).

Lastly, we revisited the mean BF state tuning curves, this time stratifying responses by both modulation motif and excitatory subtype (**Figure 3F**, **Figure 3G**). For cells assigned to the increasing motif, response magnitude scaled with pupil size across all subtypes, but the strength and pattern of this modulation varied significantly (**Figure 3G**, two-way ANOVA, *n* = 759, main effect for pupil and cell type, *p <* 10^−100^ and *p* = 5.18 × 10^−6^, respectively; cell type × pupil interaction, *p* = 3.77 × 10^−22^). A similar pattern was observed for the decreasing motif, where responses weakened with larger pupil diameter, in a subtype-dependent manner (**Figure 3G**, two-way ANOVA, *n* = 443, main effect for pupil and cell type, *p <* 10^−100^ and *p* = 0.017, respectively; cell type × pupil interaction, *p* = 2.88 × 10^−7^). For the inverted-U motif, response magnitude varied significantly with pupil size and differed by excitatory subtype (**Figure 3G**, *n* = 670, two-way ANOVA, main effect for pupil and cell type, *p <* 10^−100^ and *p* = 0.0159, respectively; cell type × pupil interaction, *p* = 1.51 × 10^−18^). In contrast, neurons exhibiting a U-shaped motif showed significant modulation by pupil diameter overall, but no systematic differences between subtype (**Figure 3G**, n = 224, two-way ANOVA, main effect for pupil and cell type, *p* = 9.16 × 10^−54^ and *p* = 0.3169, respectively; cell type × pupil interaction, *p* = 0.9203).

When responses were averaged across all motifs, pupil-linked modulation appeared relatively weak (**Figure 3A**). However, parsing by motif revealed robust and statistically significant state-dependent effects that would otherwise be obscured by population averaging (**Figure 3G**). For example, although 18–25% of cells in each subtype decreased their evoked activity with increasing pupil size (**Figure 3E**), this suppressive trend was masked in the pooled data, likely due to cancellation by coexisting increasing responses. To test whether this modulation was specific to responses at the best frequency, we repeated the clustering analysis using responses to each neuron’s least-preferred tone (“worst frequency”). This revealed the same pupil-state modulation motifs, as well as a non-modulated motif, which again varied significantly across cell types (**Supplementary Figure 2H–J**, chi-squared test, *p <* 10^−100^ and *p* = 1.9 × 10^−5^, respectively). These results indicate that pupil-linked modulation is a general feature of ACtx excitatory neurons, not restricted to responses at preferred stimuli.

### Pupil-linked arousal differentially shapes tuning selectivity

Given that response magnitudes at BF varied with arousal, we hypothesized that frequency tuning curves would also shift across pupil states. To test this, we split trials into low and high arousal states and computed separate frequency tuning curves for each neuron in each state. We then performed a linear regression by fitting each neuron’s high arousal tuning curve as a function of its low-arousal tuning curve, allowing us to quantify state-dependent transformations in tuning selectivity (**Figure 4A**). This approach yields two key parameters: the slope and y-intercept of the fit. Slopes significantly different from +1 reflect changes in tuning gain—specifically, values *<* 1 indicate divisive transformations, and values *>* 1 indicate multiplicative transformations. Likewise, y-intercepts significantly different from 0 reflect additive (positive) or subtractive (negative) shifts in overall response magnitude. The magnitude of these transformations increases with the distance from slope = 1 and intercept = 0. In many cases, both gain and offset changes occurred concurrently, reflecting compound transformations such as multiplicative/additive (**Figure 4B**) or divisive/subtractive (**Figure 4C**) effects.

**Figure 4:**
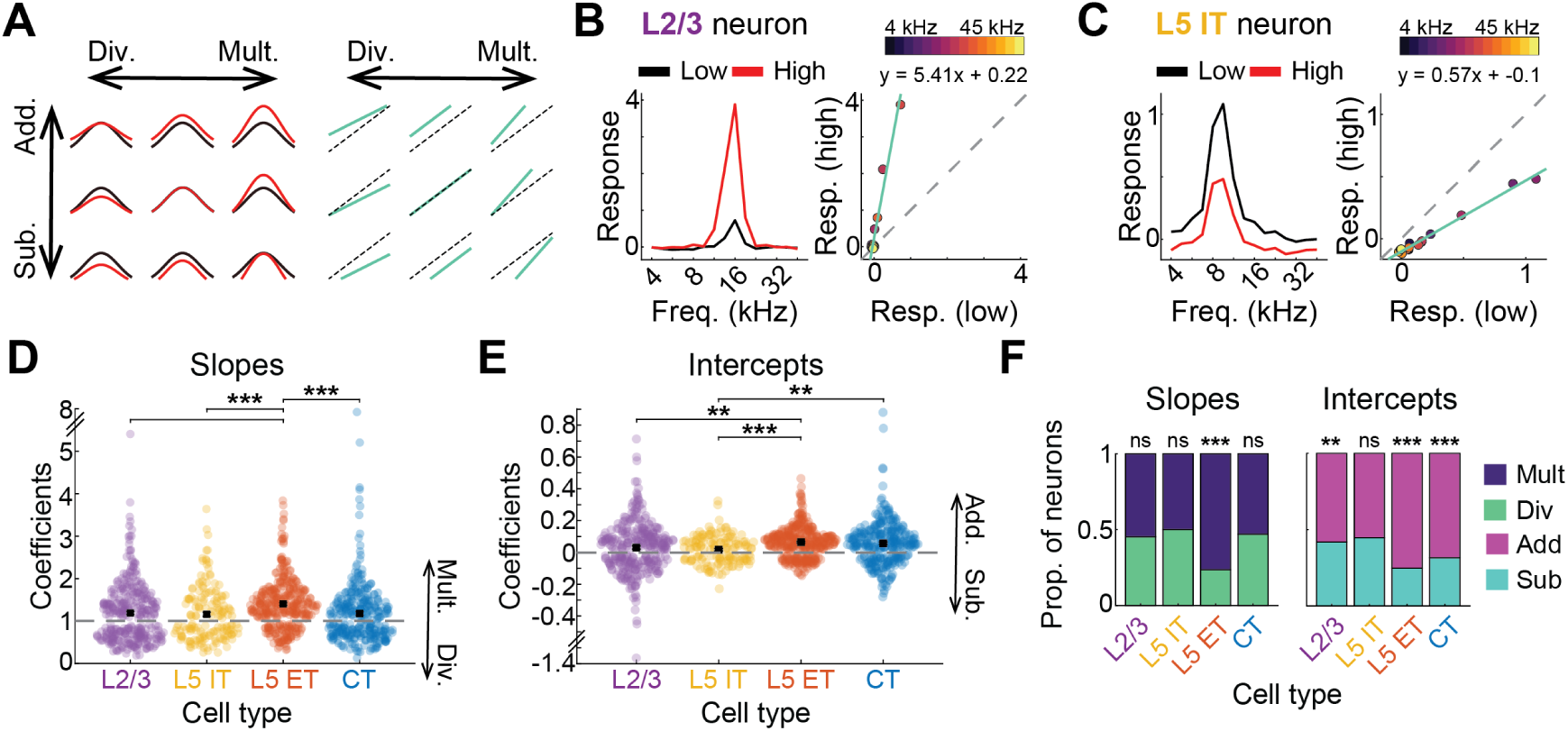
Pupil-linked arousal differentially shape tuning selectivity. **A**: Schematic illustrating how tuning curves shift from low (black) to high (red) pupil-linked arousal (left). Corresponding linear fits (turquoise) comparing high-to low-arousal tuning, overlaid with a dashed unity reference line (right). Transformation types: Add = additive, sub = subtractive, mult = multiplicative, div = divisive. **B**: Example of a multiplicative and additive transformation in a single L2/3 neuron. Frequency tuning curves at low (black) and high (red) arousal states (left). Scatter plot and linear fit (turquoise) showing the relationship between low- and high-arousal responses (right). Equation shown above. Low and high arousal is defined as pupil bins of 0 to 52% and 72% to 100% max dilation, respectively. **C**: Same as (B), for a subtractive and divisive transformation in an example L5 IT neuron. **D**: Swarm plot of significant slope coefficients across cell types, representing multiplicative/divisive transformations. Each point is a neuron (*n* = 1209, Kruskal–Wallis test, *χ*^2^ = 51.19, *p* = 4.46 × 10^−11^). Post-hoc comparisons represented as: *** *p <* 0.0001. **E**: Same as in (D), for y-intercept coefficients corresponding to additive and subtractive transformations (*n* = 1105, Kruskal–Wallis test, *χ*^2^ = 25.75, *p* = 1.08 × 10^−5^). Post-hoc comparisons represented as: ** *p <* 0.01, *** *p <* 0.0001. **F**: Proportion of neurons exhibiting multiplicative/divisive (left) or additive/subtractive (right) tuning transformations. Distributions differed significantly across cell types for multiplicative/divisive (chi-square test of independence, *χ*^2^ = 58.59, *p* = 1.17×10^−12^), and additive/subtractive transformations (chi-square test of independence, *χ*^2^ = 29.85, *p* = 1.48×10^−6^). Binomial tests were conducted for each cell type and transformation pair; ns = not significant, * *p <* 0.05, *** *p <* 0.0001.

We observed a range of state-dependent transformations in tuning gain and offset that varied significantly across excitatory subtypes. Slope coefficients, which reflect multiplicative or divisive changes, differed across groups (**Figure 4D**, *n* = 1209, Kruskal–Wallis test, *p* = 4.46×10^−11^). Post hoc comparisons revealed that L5 ET neurons had significantly different multiplicative/divisive effects compared to all other subtypes (Dunn’s test; L2/3, *p* = 5.19 × 10^−8^; L5 IT, *p* = 6.32 × 10^−6^; CT, *p* = 9.72 × 10^−9^), whereas L2/3, L5 IT, and CT neurons showed statistically indistinguishable distributions (Dunn’s test, *p >* 0.99). Similarly, y-intercept coefficients, reflecting additive or subtractive transformations, also varied by subtype (**Figure 4E**, *n* = 1105, Kruskal–Wallis test, *p* = 1.08 × 10^−5^). L5 ET neurons again differed significantly from both L2/3 and L5 IT neurons (Dunn’s test, *p* = 0.0026 and *p* = 5.74 × 10^−5^, respectively), and intercept values for CT neurons differed significantly from those of L5 IT neurons (Dunn’s test, *p* = 0.0033). These subtype-specific differences remained evident when examining the proportion of neurons classified as multiplicative/divisive or additive/subtractive (**Figure 4F**, chi-squared test, *p* = 1.17 × 10^−12^ and *p* = 1.48 × 10^−6^, respectively). To further test whether specific subtypes were biased toward particular transformation types, we performed binomial tests within each group. From low to high arousal, L5 ET neurons showed a strong preference for multiplicative gain modulation (*p* = 5.99 × 10^−25^), a pattern not observed in L2/3, L5 IT, or CT neurons (*p* = 0.08, *p >* 0.99, *p* = 0.29, respectively). L5 ET neurons also showed a significant bias toward additive shifts at higher arousal (*p* = 4.31 × 10^−22^), a trend that was also present in L2/3 (*p* = 0.0045) and CT (*p* = 1.7 × 10^−10^) neurons, but not in L5 IT neurons (*p* = 0.208). Taken together, these results indicate that L5 ET neurons exhibit the strongest and most consistent arousal-dependent transformations, characterized by increases in both gain and offset. This pattern is consistent with decreased frequency selectivity and enhanced evoked responses under high-arousal conditions.

### Pupil-linked arousal differentially modulates correlated activity

Fluctuations in pupil size have been linked to changes in the functional organization of cortical networks, including shifts in interneuronal correlations that accompany different arousal states [11, 35, 36, 56]. Yet, how neural activity co-varies within defined excitatory subpopulations remains poorly understood. Prior studies have reported that pairwise neuronal correlations tend to decrease with increasing inter-somatic distance, but it is unclear whether this spatial dependence generalizes across distinct excitatory subtypes [57–63]. To address these questions, we analyzed pure tone responses, examining the relationship between pupil-linked arousal, inter-somatic distance, and pairwise correlations. We focused on two widely used measures of correlated activity: noise correlations, which reflect shared trial-to-trial variability, and signal correlations, which capture the similarity of stimulus tuning across neuron pairs [57, 64–67] (**Figure 5A-B**).

**Figure 5:**
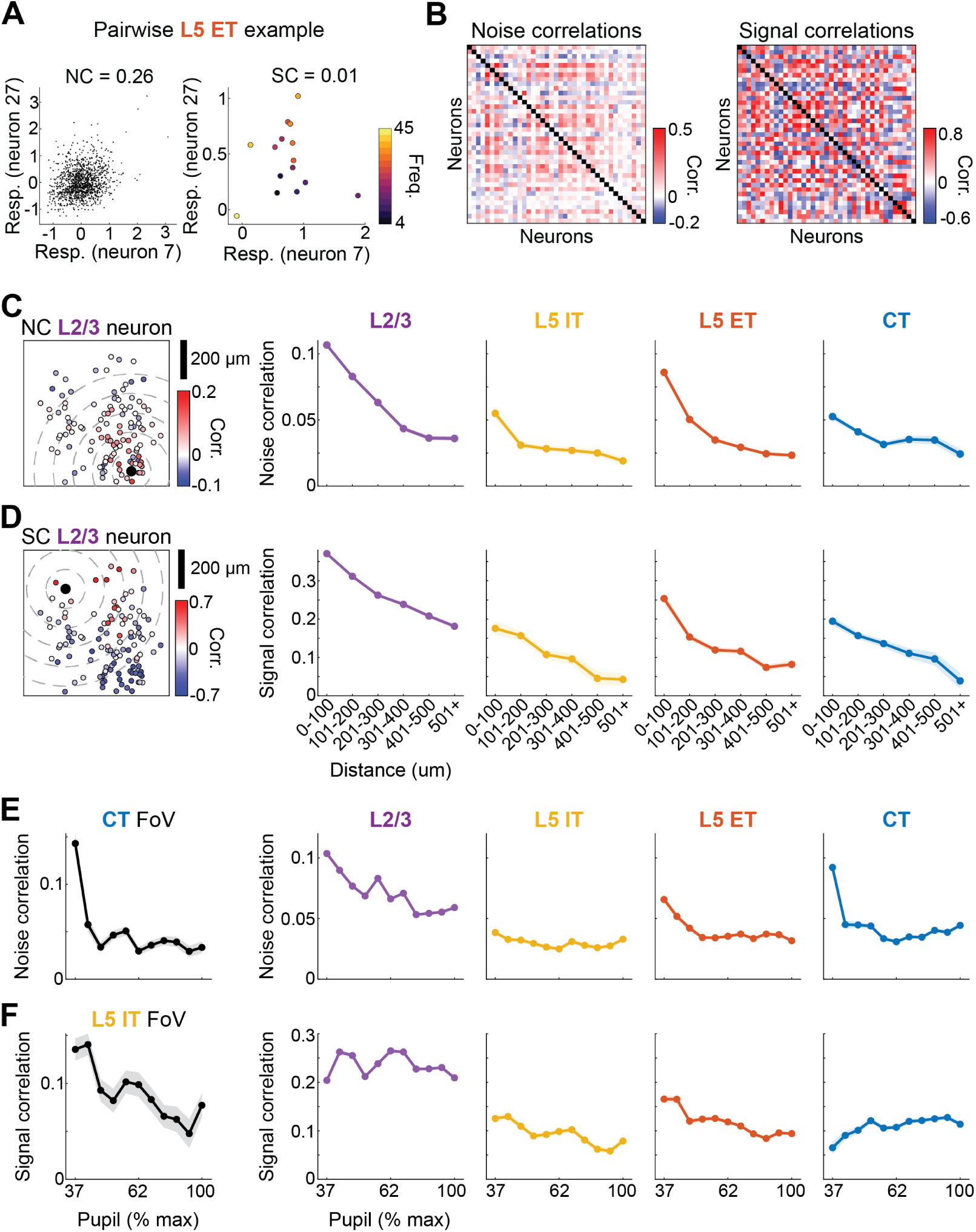
Pupil-linked arousal differentially modulates correlated activity. **A**: Example noise correlation (left) and signal correlation (right) from a pair of L5 ET neurons from the same FoV in (B). Each dot in the noise correlation plot represents the mean z-scored response on a single trial; each dot in the signal correlation plot represents the mean response to a single stimulus. **B**: Full pairwise correlation matrices for noise (left) and signal (right) correlations in an example L5 ET FoV. **C**:Pairwise noise correlations (colored dots) plotted as a function of inter-somatic distance for one example L2/3 neuron (black dot) within a single FoV (left). Mean noise correlations binned by distance for each cell type (right). Noise correlations significantly differed across distance and cell type (two-way mixed effects model, main effect for distance and cell type, *F* = 164.74 and 577.13, *p <* 10^−100^ and *p <* 10^−100^, respectively; cell type × distance interaction term, *F* = 9.82, *p* = 8.78 × 10^−24^). Pairwise cell counts: L2/3: *n* = 47, 285; L5 IT: *n* = 4, 720; L5 ET: *n* = 16, 015; CT: *n* = 8, 105). Shaded regions denote mean ± s.e.m. **D**: Same as (C), but for signal correlations. Same example FoV in (C) is used (left). Signal correlations significantly differed across distance and cell type (two-way mixed effects model, main effect for distance and cell type, *F* = 88.82 and 707.94, *p* = 1.75 × 10^−93^ and *p <* 10^−100^, respectively; cell type × distance interaction term, *F* = 3.74, *p* = 1.16 × 10^−6^). **E**: Mean noise correlations across pupil states for an example CT FoV (left) and pooled across all populations (right). Noise correlations significantly differed across distance and cell type (two-way mixed effects model, main effect for pupil and cell type, *F* = 109.33 and 2579.09, *p <* 10^−100^ and *p <* 10^−100^, respectively; cell type × pupil interaction term, F = 29.78, *p <* 10^−100^). Shaded regions denote mean ± s.e.m. **F**: Same as in (E), but for signal correlations, using an example L5 IT FoV (left). Signal correlations significantly differed across distance and cell type (two-way mixed effects model, main effect for pupil and cell type, *F* = 17.36 and 4726.36, *p* = 5.09 × 10^−32^ and *p <* 10^−100^, respectively; cell type × pupil interaction term, *F* = 49.49, *p <* 10^−100^).

We found that mean noise correlations decreased as a function of inter-somatic distance across all excitatory subtypes, regardless of pupil state (**Figure 5C**). Although this distance-dependent decrease was consistent across groups, the overall magnitude of noise correlations differed significantly by subtype (**Figure 5C**, two-way mixed-effects model, main effect for distance and cell type, *p <* 10^−100^ and *p <* 10^−100^, respectively; distance x cell type interaction, *p* = 8.78 × 10^−24^). Signal correlations also decreased with distance (**Figure 5D**) and varied significantly across subtypes (two-way mixed-effects model, main effect for distance and cell type, *p* = 1.75 × 10^−93^ and *p <* 10^−100^, respectively; distance x cell type interaction, *p* = 1.16 × 10^−6^).

Previous studies have shown that noise correlations in L2/3 neurons decrease with pupil dilation, reflecting reduced shared variability during heightened arousal [11, 35, 36, 56]. In contrast, findings for signal correlations in L2/3 have been inconsistent, with reports of both increases and decreases across arousal states [11, 35]. Here, we found that noise correlations were significantly modulated by both pupil state and excitatory subtype (**Figure 5E**, two-way mixed-effects model, main effect for pupil and cell type, *p <* 10^−100^ and *p <* 10^−100^, respectively; pupil x cell type interaction, *p <* 10^−100^). Specifically, noise correlations decreased at higher arousal levels across all excitatory subtypes, not just in L2/3, suggesting that reduced shared variability is a general feature of arousal-related cortical dynamics. In contrast, signal correlations showed a more heterogeneous pattern across subtypes, being significantly influenced by both pupil state and excitatory identity (**Figure 5F**, two-way mixed-effects model, main effect for pupil and cell type, *p* = 5.09 × 10^−32^ and *p <* 10^−100^, respectively; pupil x cell type interaction, *p <* 10^−100^). In L5 IT and ET neurons, signal correlations decreased with increasing arousal, whereas CT cells exhibited the opposite trend, with signal correlations increasing at higher pupil states. In L2/3, signal correlations were highly variable and showed no clear direction of change. These divergent effects suggest that arousal can either increase the independence of stimulus representations (as in L5 IT and ET cells) or enhance their overlap (as in CT cells), depending on the projection class.

Together, these results indicate that both arousal state and excitatory cell identity influence the structure of pairwise correlations. Specifically, pupil-linked arousal reduces shared trial-by-trial variability across the network and differentially modulates representational similarity within distinct excitatory subtypes.

### Pupil-linked arousal differentially shapes stimulus decoding performance

We have shown that pupil-linked state dynamics modulate stimulus representations at the single-cell level. Such modulation may translate to differences in how accurately sound identity is encoded across arousal states. To test this, we trained an artificial neural network classifier to decode stimulus identity from population-level neural activity, irrespective of sound-responsiveness, across multiple pupil states and excitatory subtypes. We found that decoding accuracy varied significantly with both arousal and cell type (**Figure 6A**, two-way mixed-effects model, main effect for pupil and cell type, *p* = 6.72 × 10^−5^ and *p* = 9.94 × 10^−5^, respectively; pupil x cell type interaction, *p* = 0.0018). In L2/3 and CT neurons, decoding accuracy followed an inverted-U profile across arousal states, whereas L5 IT neurons showed stable performance and L5 ET neurons exhibited improved decoding at higher arousal. Normalizing decoding performance within each FoV to its peak further confirmed this cell-type-specific effect (**Figure 6B**, mixed-effects model, main effect for pupil and cell type, *p* = 8.13 × 10^−10^ and *p* = 0.028, respectively; pupil x cell type interaction, *p* = 0.057).

**Figure 6:**
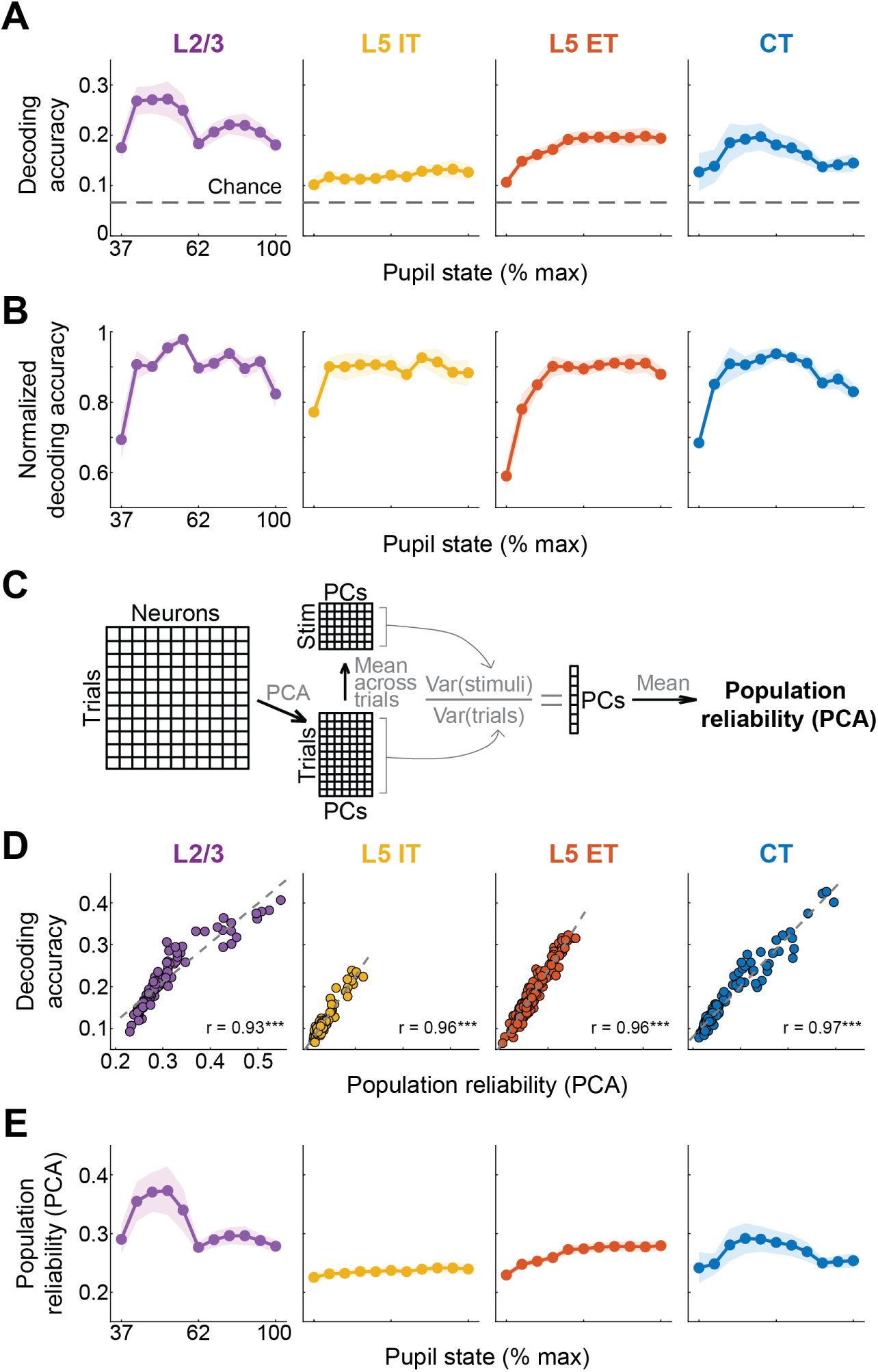
Pupil-linked arousal differentially shapes stimulus decoding performance. **A**: Mean stimulus decoding accuracy across pupil states for each excitatory cell type. Decoding performance was significantly influenced by pupil size and cell type (two-way mixed-effects model, main effect for pupil and cell type, *F* = 11.49 and 7.91, *p* = 6.72 × 10^−5^ and *p* = 9.94 × 10^−5^, respectively; pupil x cell type interaction term, *F* = 1.995, *p* = 0.0018). Chance accuracy is 1/15. Shaded regions denote mean ± s.e.m. **B**: Same as in (A), but decoding accuracy is normalized within each FoV to its maximum value to yield a proportion of peak classification performance (value of 1). Normalized decoding accuracy differed across pupil states and cell type (two-way mixed effects model, main effect for pupil and cell type, *F* = 13.07 and 3.18, *p* = 8.13 × 10^−10^ and *p* = 0.028, respectively; cell type × pupil interaction term, *F* = 1.47, *p* = 0.057). Shaded regions denote mean ± s.e.m. **C**: Schematic depicting the computation of PCA-derived population reliability, defined as the proportion of trial-averaged variance explained relative to total trial-to-trial variability. **D**: Scatter plot showing the relationship between mean stimulus decoding accuracy and PCA-derived population reliability for each FoV across pupil states and imaging days. Decoding performance and population reliability were highly correlated for every cell type (L2/3, *r* = 0.93,*p* = 6.29 × 10^−42^; L5 IT, *r* = 0.96, *p* = 1.07 × 10^−57^; L5 ET, *r* = 0.96, *p* = 7.22 × 10^−96^; CT, *r* = 0.97, *p* = 7.67 × 10^−74^). **E**: Mean PCA-derived population reliability across pupil states. Reliability significantly varied as a function of both pupil size and cell type (two-way mixed effects model, main effect for pupil and cell type, *F* = 9.98 and 6.74, *p* = 0.0002 and *p* = 0.0004, respectively; cell type × pupil interaction term, *F* = 2.19, *p* = 0.0004). Shaded regions denote mean ± s.e.m.

We next examined whether network-level features influence decoding performance, focusing first on noise correlations, which can limit the amount of independent information encoded by a population [66, 68–70]. However, the functional consequences of such correlations remain contentious, with some studies reporting little impact on population coding fidelity [71]. To directly test their contribution, we shuffled trials within each stimulus class for each neuron/frequency pair, thereby disrupting noise correlations while preserving mean responses. Eliminating shared variability had no measurable effect on decoding accuracy in any excitatory subtype (**Supplementary Figure 3A-B**, mixed effects model, main effect of shuffling, *p >* 0.99 in both figures).

We next evaluated whether decoding accuracy is shaped by population reliability, the consistency of neural responses across repeated stimulus presentations, which is thought to enhance sensory decoding by stabilizing population codes that improve pattern separation [41]. To assess this, we quantified population-level reliability across FoVs, pupil states, and recording sessions, and compared these values to decoding performance within each excitatory cell type. Reliability was estimated using principal component analysis (PCA): for each principal component, we computed the ratio of the variance of the mean stimulus-evoked response to the total trial-to-trial variance, and then averaged across components to yield a single reliability score (**Figure 6C**). This measure reflects the stability of evoked population activity patterns, with higher values reflecting greater reliability. Across all excitatory subtypes, we observed strong positive correlations between decoding accuracy and population-level reliability (**Figure 6D**; L2/3, *p* = 6.29 × 10^−42^; L5 IT, *p* = 1.07 × 10^−57^; L5 ET, *p* = 7.22 × 10^−96^; CT, *p* = 7.67 × 10^−74^).

To complement our PCA-based measure, we also assessed single-neuron reliability, calculated independently for each sound-responsive neuron across all pupil states [41]. This analysis revealed significant differences in reliability across excitatory subtypes, independent of pupil state, with L5 ET neurons exhibiting the highest and L5 IT neurons the lowest reliability (**Supplementary Figure 3C**, Kruskal–Wallis test, *p* = 2.7 × 10^−9^). PCA-derived and single-neuron estimates of population reliability were highly correlated within each cell-type (**Supplementary Figure 3D**; L2/3, *p* = 1.7 × 10^−33^; L5 IT, *p* = 7.89 × 10^−83^; L5 ET, *p* = 2.72 × 10^−98^; CT, *p* = 2.01 × 10^−62^). Moreover, like PCA-derived reliability, single-neuron reliability was strongly predictive of decoding accuracy across all excitatory populations (**Supplementary Figure 3E**; L2/3, *p* = 4.53 × 10^−34^; L5 IT, *p* = 1 × 10^−51^; L5 ET, *p* = 8.97 × 10^−93^; CT, *p* = 2.24 × 10^−70^).

Arousal-dependent changes in decoding performance likely reflect shifts in neural network dynamics. Given the tight coupling between reliability and decoding accuracy, we next asked whether reliability itself is modulated by arousal, and could serve as a mechanistic link between internal state and sensory encoding. To test this, we computed mean reliability within each pupil-state bin and found that PCA-derived population-level reliability varied significantly with arousal in a cell-type-specific manner (**Figure 6E**, two-way mixed-effects model, main effect for pupil and cell type, *p* = 0.0002 and *p* = 0.0002, respectively; pupil x cell type interaction, *p* = 0.0004). Single-neuron reliability showed the same pattern (**Supplementary Figure 3F**, two-way mixed-effects model, main effect for pupil and cell type, *p* = 5.17 × 10^−5^ and *p* = 2.67 × 10^−8^, respectively; pupil x cell type interaction, *p* = 0.0012). Across both metrics, reliability tracked decoding performance across pupil states: L2/3 and CT neurons showed inverted-U profiles, L5 IT neurons remained largely unchanged, and L5 ET exhibited a monotonic increase in reliability with arousal (**Figure 6A** and **6E; Supplementary Figure 3F**). These results demonstrate that the fidelity of sound representations is shaped by both cell type and internal state. Critically, the stability of the population code emerges as a key constraint on how effectively sensory stimuli are decoded from cortical activity.

## Discussion

We used pupil diameter as a proxy for arousal to investigate how internal state shapes sensory processing in the neocortex. Through simultaneous two-photon calcium imaging and pupillometry in awake mice, we systematically characterized arousal-dependent changes in response magnitude, frequency tuning, interneuronal correlations, and stimulus decoding across major excitatory subpopulations in the ACtx: intratelencephalic (IT), extratelencephalic (ET), and corticothalamic (CT) neurons. By employing a cell-type-specific approach, our study addresses longstanding discrepancies in the literature concerning how pupil-linked arousal affects sensory cortex activity. We provide evidence that these divergent findings can be reconciled by considering the distinct, subtype-specific ways in which arousal modulates neural activity. First, we identified a rich diversity of arousal-dependent response motifs, including both linear and non-linear patterns, that were unevenly distributed across subtypes and masked in population-averaged analyses (**Figure 3**). Second, we predominantly found multiplicative and additive changes in frequency tuning curves at higher arousal levels specifically in L5 ET neurons, whereas other populations exhibited weaker and more variable changes in tuning (**Figure 4**). Third, trial-to-trial variability (noise correlations) decreased with increasing arousal across all groups, but representational similarity (signal correlations) showed divergent, cell-type-specific trends: decreasing in L5 IT and ET neurons, and highly variable in L2/3 (**Figure 5**). Finally, decoding accuracy varied systematically with arousal in a cell-type-specific manner, driven in part by differences in population-level response reliability (**Figure 6**). Together, these findings show that the influence of pupil-linked arousal on sensory coding is not homogeneous across the cortex but instead arises through distinct mechanisms in different excitatory subpopulations, highlighting the need for cell-type specific resolution when interpreting state-dependent neural dynamics.

### Effects of neuromodulation in the neocortex

Global arousal states are regulated by a network of subcortical neuromodulatory systems that broadly influence brain and behavior. Many of these neuromodulatory systems, including the locus coeruleus (norepinephrine: LC-NE), basal forebrain (acetylcholine: BF-ACh), and dorsal raphe (serotonin: DR-5-HT), correlate with fluctuations in pupil diameter and innervate all layers of sensory cortex, where they release neuromodulators that alter cortical dynamics [12, 20–26, 28–33]. These neuromodulators modulate spontaneous cortical activity and modulate evoked responses by dynamically shifting cortical excitability. Moment-to-moment fluctuations in neuromodulatory tone, indexed by pupil size, can therefore alter how sensory inputs are processed. Such modulation has been shown to affect receptive field properties and tuning selectivity, as demonstrated in visual and auditory cortices following cholinergic agonist application or basal forebrain stimulation [32, 72–76]. Similar effects have been observed for LC-NE and DR-5-HT systems [77–83].

Several mechanisms may underlie the cell-type-specific effects observed in our study. First, neuromodulatory afferents from the BF, LC, and DR exhibit laminar specificity, with axon terminal density varying across cortical layers [84–88]. Second, receptor expression for ACh, NE, and 5-HT varies across cortical layer and by excitatory cell type [86, 89–91]. Third, electrophysiological studies have shown that excitatory subtypes respond differently to the same neuromodulators. In L6, for example, CT and ET neurons exhibit ACh-mediated depolarization, whereas L6a and L6b IT neurons show either hyperpolarization or depolarization depending on the subtype [92]. In L5, serotonin excites IT neurons but inhibits ET neurons via distinct 5-HT receptor types [93–95]. Moreover, both NE and ACh (via *α*2-adrenergic and muscarinic receptors, respectively) preferentially increase the excitability and persistent firing of L5 ET neurons compared to L5 IT neurons [27, 96, 97]. This *in vitro* evidence is consistent with our *in vivo* finding that, across pupil-linked arousal states, L5 ET neurons increase their neuronal gain relative to L5 IT neurons—a pattern that aligns with the known coupling between pupil size and BF-ACh and LC-NE terminal activity in sensory cortex [12]. Optogenetic activation of cholinergic terminals in visual cortex, mimicking heightened arousal, preferentially induces multiplicative and additive gain effects in L5 compared to L2/3 cells [76]. Assuming similar ET/IT responsiveness across auditory and visual cortices would suggest that this effect is most prominent in L5 ET cells.

Arousal-dependent reductions in correlated activity may also be mediated by neuromodulators. In visual cortex, BF stimulation alters firing rates across layers but consistently reduces correlated variability across neurons [98]. This observation aligns with our finding that noise correlations decreased with increasing arousal across all excitatory subpopulations, a single-cell and cell-type-specific extension to corollary EEG and LFP studies showing cortical desynchronization under heightened arousal (**Figure 5**) [4, 99].

Importantly, no single neuromodulator alone can account for the full range of cell-type-specific effects we observed. Arousal-associated changes in brain state likely reflect a multiplexed neuro-modulatory mélange that alters cortical dynamics acting through both synaptic and volume transmission mechanisms. Subcortical neuromodulatory neurons can co-release glutamate and GABA in addition to their primary neuromodulator [100–103], and LC terminals have been shown to release both NE and dopamine, which differentially modulate excitatory cell response [92, 103, 104]. Astrocytes also respond to neuromodulators and may further shape state-dependent sensory processing [105]. The observation that not all neurons of a given subtype exhibit the same arousal-linked modulation (e.g., L5 ET neurons with differing pupil-state tuning motifs) likely reflects heterogeneity in presynaptic input patterns, receptor localization, and downstream intracellular signaling. This variability underscores the need to consider not only cell type but also microcircuit context and input–output relationships when interpreting state-dependent changes in cortical function.

### Sources and significance of non-linear gain modulation

While arousal promotes alertness and facilitates stimulus detection, heightened arousal can lead to impulsivity and impair performance on demanding tasks [9, 20, 106, 107]. Conversely, low arousal is associated with disengagement and diminished behavioral responsiveness. The Yerkes–Dodson law, as formalized by Hebb, posits an inverted-U relationship between arousal and behavioral performance, whereby moderate arousal levels yield optimal outcomes, while both hypo- and hyper-arousal degrade performance [108, 109]. This curvilinear relationship has been widely supported across species, sensory modalities, and behavioral paradigms [9, 38, 99, 106, 110–117]. Despite its broad behavioral relevance, direct neural correlates of this relationship have remained elusive.

We observed both linear and non-linear relationships between arousal and evoked response magnitude. Linear effects, especially monotonic increases with pupil size, were the most prevalent, consistent with the broader literature on pupil-linked neural activity [11, 35–39]. While some studies emphasize a non-monotonic but not strictly quadratic function [118], an inverted-U pattern, mirroring the Yerkes-Dodson relationship, is commonly reported and favored as a unifying principle [9, 36, 38]. For example, MEG in humans shows that the spectral power of cortical activity follows distinct arousal-related trends across frequency bands, with an inverted-U pattern observed specifically in the 8–16 Hz range [38]. In ferret ACtx, arousal-related gain modulation motifs include increasing, decreasing, and both positive and negative quadratic patterns [36]. Our results confirm the presence of such non-linear tuning across cortical excitatory populations, while also reinforcing the predominant linear trends found in prior work (**Figure 3**). Critically, we show that the distribution of arousal modulation motifs is not uniform across excitatory subtypes. L5 ET and IT populations differed by approximately 10% in the relative proportion of increasing versus inverted-U tuning motifs. L5 ET cells had the highest proportion of increasing-modulation cells and the lowest proportion of inverted-U cells, whereas the opposite pattern was more pronounced in L5 IT and CT populations. These findings suggest that excitatory cell subtypes possess distinct propensities for linear versus non-linear arousal-related gain modulation, contributing to divergent effects of brain state on cortical processing.

The mechanistic origin of these state-dependent non-linear shifts in cortical activity remains uncertain. One possibility is that they are inherited from subcortical sources. Prior work has shown that multiunit responses in the auditory thalamus exhibit peak responsiveness at intermediate arousal levels, suggesting a feedforward source of non-linear modulation [9]. Given the canonical thalamocortical circuit, such modulation could be relayed to layer 4 (the principal cortical recipient of thalamic input), then to L2/3, and ultimately to L5 [119, 120]. CT neurons, in particular, are well positioned to receive such modulation, as their apical dendrites frequently arborize within L4 [51]. At the same time, CT neurons may shape thalamic activity through their corticothalamic projections, potentially supporting a recurrent loop wherein cortical and thalamic dynamics coregulate state-dependent activity patterns.

Alternatively, intracortical mechanisms may contribute to non-linear gain modulation. One proposed model implicates the interaction between vasoactive intestinal peptide-expressing (VIP) and somatostatin-expressing (SOM) inhibitory interneurons [116]. As arousal increases, VIP neurons suppress SOM neurons, disinhibiting excitatory cells and increasing gain. At high arousal, VIP activity plateaus, allowing SOM-mediated inhibition to reduce excitatory drive, resulting in an inverted-U shaped pattern. This model is supported by observations that VIP and SOM cells tend to exhibit opposing pupil-linked activity patterns, with VIP activity strongly tracking pupil size, while correlations between pupil size and SOM activity are more heterogeneous [11, 42]. Future experiments are needed to directly test these candidate mechanisms. Specifically, dissecting the relative contributions of feedforward thalamic input and intracortical inhibitory circuitry may help clarify how non-linear, state-dependent modulation arises within specific excitatory subtypes.

### Influences on neural discriminability

Our results demonstrate that sound encoding and decoding vary with pupil-linked arousal in a cell-type-specific manner. While decoding performance in L5 IT neurons remained largely stable across arousal states, L2/3 and CT populations exhibited an inverted-U relationship, and L5 ET neurons showed enhanced decoding accuracy at higher arousal levels. The inverted-U trend in L2/3 aligns with recent findings suggesting that intermediate arousal states facilitate stimulus decoding [40]. In contrast, the stable decoding performance of L5 IT neurons may reflect a role in signal detection that is relatively invariant to changes in brain state. Although L5 IT neurons had the lowest overall decoding performance and weakest average sound responsiveness, their activity nonetheless conveyed identity-relevant information, indicating latent population-level structure. By comparison, the progressive improvement in decoding accuracy in L5 ET neurons with increasing arousal supports the idea that heightened brain states enhance sensory representations to facilitate environmental awareness.

Reliable population codes, characterized by low trial-to-trial variability, are thought to support more accurate decoding of sensory stimuli [41, 98]. Consistent with this, we found strong positive correlations between decoding performance and population-level reliability across all cell types (**Figure 6**). Moreover, the pupil-dependent reliability profiles closely mirrored the corresponding decoding trends: inverted-U shaped for L2/3 and CT, flat for L5 IT, and monotonically increasing for L5 ET neurons. These findings suggest that shifts in neural response reliability across arousal states exert a potent influence in modulating the fidelity of sensory representations across cortical excitatory subtypes.

### Limitations and considerations

Arousal state is closely linked to locomotion, which dilates the pupil and can suppress ACtx activity via recruitment of local inhibitory circuits [9, 11, 56, 121, 122]. Although we did not explicitly control for movement, mice were head-fixed on a stationary platform and extensively habituated prior to imaging, which minimized spontaneous movement during recordings. Critically, elevated arousal can occur independent of movement, and our clustering analysis revealed a broad diversity of arousal-linked modulation motifs. This heterogeneity suggests that any residual movement-related effects likely averaged out across the dataset.

The pupil-dependent effects observed within a given cell type may reflect not only direct neuromodulatory influences, but also indirect consequences of intracortical circuit dynamics. For example, activation of CT neurons in both visual and auditory cortices can alter activity across the cortical laminae [48, 49, 51]. Such interactions imply that the neuromodulator-driven activation of one subpopulation, such as CT cells, could propagate effects to other subtypes through local connectivity. Disentangling these global versus local contributions will require future experiments that combine cell-type-specific manipulation of neuromodulatory inputs with simultaneous recordings across multiple cortical layers.

Finally, the generalizability of our findings across cortical areas and behavioral contexts remains to be established. Prior studies have shown that the effects of pupil-linked arousal vary by brain region and are sensitive to behavioral engagement, task demands, and stimulus complexity [6, 110, 117, 123–126]. Our study was limited to passive listening with pure tones, and future work should test whether similar patterns of state-dependent modulation are observed under more ethologically relevant conditions, including naturalistic sounds and active task engagement.

## Conclusions

Arousal systems play a critical role in adaptive behavior by modulating sensory processing in accordance with internal state. Our findings demonstrate that distinct excitatory subpopulations within ACtx exhibit unique patterns of arousal-dependent modulation, influencing response magnitude, tuning selectivity, correlated activity, and stimulus encoding. This cell-type-specific diversity in state-dependent dynamics likely underpins the brain’s ability to flexibly encode sensory information under varying arousal conditions. Moreover, these distinctions offer a unifying framework to reconcile previously conflicting reports on how arousal shapes sensory processing. Together, our results underscore the importance of cell-type-specific approaches in understanding the neural basis of state-dependent perception and cognition.

## Materials and Methods

### Mice

All procedures were approved by the University of Pittsburgh Animal Care and Use Committee and followed the guidelines established by the National Institute of Health for the care and use of laboratory animals. We expressed GCaMP8s in adult mice of both sexes for all two-photon calcium imaging experiments. The mouse strains used were: C57BL/6 (L2/3, *N* = 6; L5 ET, *N* = 10), Tlx3-PL56-Cre (*N* = 7), and Ntsr1-Cre (*N* = 9). All mice were light-reversed (12 h dark/light cycle) with ad libitum access to food and water throughout experiments. All imaging was conducted during the dark cycle.

### Surgical Procedures

All surgical procedures were conducted in anesthetized mice under aseptic conditions. Mice were anesthetized with 5% isoflurane in oxygen and maintained at 1-2% throughout surgery. Lidocaine hydrochloride (local anesthetic) was injected subcutaneously prior to any incision of the skin. Ophthalmic ointment was applied over the eyes to prevent dryness. After surgery, mice received a subcutaneous injection of carprofen (5 mg/kg) and carprofen MediGel was provided in their home cages for 3 days post-operation. All mice underwent a virus delivery and a cranial window implantation surgery.

### Virus-mediated gene delivery

All virus injections were conducted on 7-10-week-old mice. Injections for L2/3, L5 IT, and CT cohorts were delivered into the right auditory cortex at a depth of ∼500 *µ*m under the dura. A single incision was made over the right temporal ridge. Virus injections were made using pulled-glass pipettes and a programmable injector (Nanoject 3, Drummond Scientific) to deliver virus through two ∼200 *µ*m burr holes (250 nl per hole, 500 nl total). Neurons in L2/3 were targeted by injecting AAV1-syn-jGCaMP8s-WPRE (2 × 10^12^ vg/mL) into C57BL/6 mice. L5 IT and CT neurons were targeted by injecting AAV1-syn-FLEX-jGCaMP8s-WPRE (6-7 × 10^12^ vg/mL) into Tlx3-PL56-Cre and Ntsr1-Cre mice, respectively. L5 ET neurons were targeted by injecting retrograde AAV1-syn-jGCaMP8s-WPRE (7-8 × 10^12^ vg/mL) into the right inferior colliculus (IC) of C57BL/6 mice. A single incision was made along the midline, above the sagittal suture to gain access to bregma and lambda. Delivery of the virus through one burr hole at two depths (300 nl per depth, 600 nl total) was made into the IC using stereotaxic coordinates (AP, −5 mm; ML, 0.9 mm; DV, 0.8 and 0.3 mm). Viruses were diluted with PBS to acquire the desired titer and the rate of injection was 10 nl every 45-60 seconds. The glass pipette remained in the brain for an additional 10 minutes at each site following virus delivery for all injections. Incision sites were sutured and antibiotic ointment was applied.

### Cranial Window Implantation

Two weeks following virus injection, mice were anesthetized for chronic imaging window implantation surgery. An intraperitoneal injection of dexamethasone sodium phosphate (2 mg/kg) was administered to reduce inflammation and brain swelling. Skin at the top of the head was removed and an etchant was applied to the dorsal surface of the skull to facilitate head plate adhesion. A customized titanium head plate was then affixed to the skull using dental cement. A cranial window, comprising three thin glass coverslips (3-3-4mm stack), was then inserted in a 3mm diameter craniotomy over the right ACtx and secured to the skull using dental cement.

### Histology

Mice were deeply anesthetized with 5% isoflurane and transcardially perfused with and fixed in 4% paraformaldehyde in 0.01M PBS solution. Post-fixed brains were transferred to 30% sucrose the following day. Coronal sections (50 *µ*m) of the ACtx were made using a cryostat and stored in PBS. Slices were mounted onto glass slides and stained with DAPI prior to coverslipping. Fluorescence imaging was performed using an epifluorescence microscope (Thunder Imager Tissue, Leica) at 10x magnification. Channels for GFP (GCaMP8s) and DAPI were merged, color-balanced, and exported using Fiji (ImageJ).

### Calcium Imaging

Light-reversed mice were awake and head-fixed for all recording sessions. Prior to imaging, mice were habituated to head-fixation and the recording chamber for several days. Neural activity in response to four pure tones (4, 8, 16, and 32 kHz) were captured by widefield fluorescence imaging (Bergamo, ThorLabs) and used to functionally confirm the location of the right primary ACtx. Two-photon calcium imaging was conducted using an InSightX3 (Spectra Physics) Laser tuned to 940 nm and a water-immersion objective (Nikon 16x). This objective was fixed with a custom cylindrical blackout curtain to shield the PMTs from potential interference from ultraviolet (UV) light (see *Videography*). All two-photon imaging (Bergamo, ThorLabs) was of the right ACtx.

Mice were head-fixed upright with the microscope rotated to be parallel to the cranial window (approximately 40 to 50^◦^ tilt). Images were collected at 30 Hz. The depth below pial surface used for recordings depended on neuron subtype (L2/3: 150-250 *µ*m, L5 IT: 350-500 *µ*m, ET: 450-600 *µ*m, CT: 600-700 *µ*m). Separate FoVs from the same mouse were at least 50 *µ*m above or below the original imaging plane. All two-photon calcium imaging was conducted within a dark, sound-attenuating chamber so that video capture of pupil diameter was luminance-independent. Sound presentations consisted of 15 pure tone stimuli ranging from 4 kHz to ∼45.3 kHz at 0.25 octave spacing. Stimuli were generated with a 24-bit digital-to-analog converted (National Instruments model PXI-4461) using custom scripts programmed in MATLAB (Mathworks) and LabVIEW (National Instruments). Stimuli were calibrated using a wide-band ultrasonic acoustic sensor (SPM0204UD5, Knowles Acoustics). All tones were presented unilaterally to the left (contralateral) ear via a free field speaker (PUI Audio) for 50 ms and at 70 dB SPL. Pure tone stimuli were presented in a pseudo-random sequence. Trials were two seconds in duration with stimulus onset occurring at 500 ms. Mice were imaged within the same time window of their light cycle to minimize circadian variability.

### Videography

High-speed videography of the mouse’s face (left side) was recorded at 30 Hz concurrently with two-photon imaging in all sessions (Genie Nano M2020, Teledyne; TEC-55, Computar). An infrared LED provided consistent facial illumination. To prevent pupil dilation from saturating the entire eye, a small amount of UV light was used to restrict the maximum range of dilation, enabling consistent measurement of maximum pupil diameter across mice. UV light intensity was held constant across imaging days and was the sole visible light source in the recording booth.

### Image Analysis

#### Two-photon

Post-recording analysis began with Suite2p [54], an open-source image processing pipeline. It was used to register raw calcium movies, detect spatial regions of interest (ROIs), distinguish between neuronal and non-neuronal ROIs, extract calcium fluorescence signals from each ROI, and process spike deconvolution. All ROIs were then manually confirmed so that only soma activity was analyzed. To maximize the number of pupil states per FoV, a large subset of neurons were matched longitudinally across days using the open-source algorithm ROICat (https://github.com/RichieHakim/ROICaT). Neural responses were z-scored relative to average activity during a spontaneous window 500 ms preceding stimulus onset.

#### Pupil analysis

Pupil diameter was extracted from face videography recordings using DeepLabCut, an open-source deep learning-based pose estimation toolbox [127]. We trained a model on manually labeled frames to track 8 user-defined points equally distributed around the pupil perimeter. For each frame, an ellipse was fit to the pupil labels using a least-squares method and pupil size was defined as the length of the ellipse’s major axis. Marker estimates with a confidence score below 95% were excluded from the fit, and frames with fewer than 6 valid points were assigned a NaN. This thresholding approach reliably excluded eye blinks, which were confirmed to occur with low marker confidence. Biologically implausible values and sharp frame-to-frame fluctuations were removed, and the remaining trace was smoothed using a moving median filter. Missing pupil values were linearly interpolated across neighboring valid samples. Pupil diameters were normalized within each mouse to the maximum observed pupil diameter across all sessions. Pupil state for each trial was defined as the average normalized pupil diameter 500 ms prior to stimulus onset to avoid stimulus-evoked effects. Pupil was discretized into 11 states on either side of the median pupil size across all mice (**Figure 1D**, median = 62%) to produce the following bins: 0-37, 38-42, 43-47, 48-52, 53-57, 58-62, 63-67, 68-72, 73-77, 78-82, 83-100% max dilation. Changes to tuning selectivity in **Figure 4** were based on low and high pupil-linked arousal states, defined as 0-52 and 72-100% max dilation, respectively.

### Data Analysis

#### Multivariate linear regression model

We implemented a multivariate linear regression model to assess how trial-by-trial evoked neural responses are shaped by auditory and pupil state variables. The dependent variable, *y*, was the mean evoked response of a single neuron on a given trial. This response was modeled as the linear combination of stimulus identity, baseline neural activity, and baseline pupil activity using the following equation:

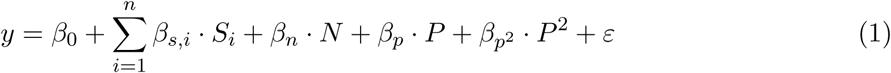

where *β*_0_ is the intercept, and the terms *S_i_* ∈ 0, 1 are a one-hot encoded representation of stimulus identity across *n* = 15 possible pure tone frequencies, each associated with a stimulus-specific coefficient *β_s,i_*. These coefficients were confirmed to align to each neuron’s frequency tuning curve by computing the Pearson correlations between *β_s_* and mean stimulus response vectors, and testing it against a null distribution generated by shuffling stimulus labels (two-sample Kolmogorov–Smirnov test, 1000 permutations, **Supplementary Figure 1A**). Baseline neural and pupil activity are denoted by *N* and *P*, with their corresponding weights *β_n_* and *β_p_*. Non-linear relationships are captured by the pupil squared term, *p*^2^, and its respective coefficient 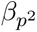 The error term *ɛ* accounts for residual variance not explained by the predictors.

Regression was performed using MATLAB’s fitrlinear function with a least-squares loss function and ridge regularization. Model coefficients (*β*) were extracted for each neuron and collated into a [neurons x coefficients] matrix. For cross-cell comparisons, the largest *β_s_* coefficient (corresponding to a neuron’s best frequency), along with *β_p_* and 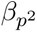 coefficients, were pooled across all neurons and z-scored independently for each coefficient type. A heuristic “sound” threshold of 0.5 standard deviations (sd) was applied to the distribution of normalized maximum *β_s_* coefficients to classify neurons as sound-responsive (*z >* 0.5 sd) or non-sound-responsive (*z* ≤ 0.5 sd). To identify neurons with the most suppressed responses (**Supplementary Figure 2H-J**), we pooled each neuron’s minimum *β_s_* coefficient (corresponding to a neuron’s worst frequency). Because all values were negative and skewed, we log-transformed their absolute values before z-scoring. Neurons exceeding a 0.5 sd threshold were classified as having strong frequency-specific suppression. Notably, sound-responsive neurons could also meet this criterion if their least-preferred frequency evoked sufficient suppression.

#### Simulating arousal modulation motifs

To simulate neural responses exhibiting distinct pupil-state modulation motifs, we generated synthetic datasets using custom MATLAB code. Five synthetic neurons were created, each tuned to the same preferred stimulus frequency but differing in how their evoked responses varied with pupil size (**Supplementary Figure 1D**). These neurons were designed to reflect one of five pupil-state modulation motifs: increasing, decreasing, inverted-U (negative quadratic), U-shaped (positive quadratic), and non-modulated (flat across pupil states). A 1200 trial design matrix was constructed using the same predictors as the multivariate linear regression model (**Supplementary Figure 1B**). Simulated baseline neural activity was sampled from a Gamma distribution and baseline pupil diameters were uniformly sampled from the interval [0,1], except for preferred stimuli trials, which evenly spanned the full [0,1] range (**Supplementary Figure 1C**, left). To simulate the response vector, a deterministic trend between pupil size and evoked neural activity was first specified for each neuron based on its assigned motif. Structured Gaussian noise was introduced to reflect trial-to-trial variability (**Supplementary Figure 1B** and **1C**, right).

#### Linear extrapolation

Missing data at the pupil-state tails (lowest and highest states) were linearly extrapolated for mean response at best frequency and hierarchical clustering analyses. Stimulus/state combinations with less than 5 trials of data were registered as NaN values and linearly interpolated and/or extrapolated, where appropriate. We devised two methods for linear extrapolation: boundless and bounded. Boundless extrapolation linearly extrapolated values at the pupil-state tail(s) without any upper or lower limits (**Supplementary Figure 2A**). Bounded extrapolation did the same but was subject to upper and lower limits that were based on the range of existing data for that neuron (**Supplementary Figure 2B**). Specifically, the maximum or minimum extrapolated value was determined by adding or subtracting 40% of the data range to the highest or lowest existing datum, respectively. Once the upper or lower limit was determined, intermediate NaN values were linearly interpolated. Solely using one of these extrapolation methods yielded some neurons with extreme response magnitudes at the tails. To address this, boundless or bounded extrapolation was applied on a neuron-by-neuron basis based on whichever extrapolation method had the smaller slope stemming from existing data. Neurons with more than 4 consecutive NaNs at either pupil state tails were not extrapolated.

#### Hierarchical clustering

To identify clusters of neurons exhibiting unique pupil-state modulation motifs, we performed hierarchical clustering on mean z-scored activity in response to a single stimulus as a function of pupil state; i.e., a neuron’s state tuning curve. Exclusively sound-responsive neurons were used for **Figure 3** and **Supplementary Figure 2C-G**, whereby mean activity was extracted from a baseline or evoked time window (500 ms pre-stimulus or 333 ms post-stimulus, respectively) using a neuron’s best frequency, defined as the stimulus that produced the largest mean activity irrespective of pupil-state. Clustering state tuning curves on neurons with the most suppressed responses (i.e., worst frequency) was based on activity during the evoked time window (**Supplementary Figure 2H-J**). At least 4 pupil states with a minimum of 5 trials per stimulus/state combination were required to obtain a neuron’s state tuning curve. Insufficient data at stimulus/state combinations were marked with NaN values and linearly interpolated and/or extrapolated, where appropriate (see *Linear extrapolation*). Neurons with one or more NaNs in their state tuning curve were excluded from this analysis, as clustering would group these together.

State tuning curves were rescaled between 0 and 1 to normalize the range of neural responses across all neurons prior to clustering. All neurons across the subtypes were pooled together and clustered using Euclidean distance and Ward’s linkage methods (MATLAB linkage and cluster functions) to extract a clustering tree. To identify a non-redundant set of clusters, we iteratively merged similar branches of the hierarchy. This process combined manual inspection with quantitative assessment using silhouette analysis (MATLAB evalclusters function), selecting the final cluster count based on the elbow of the silhouette curve to balance state-dependent diversity and parsimony. We verified that the mean z-scored and rescaled responses for each cluster were similar to ensure that minor variations in neural activity did not drive spurious clustering. Clusters were then manually grouped into one of the pupil-state modulation motifs: increasing, decreasing, inverted-U (negative quadratic), U-shaped (positive quadratic), or non-modulated. Consequently, each neuron belonged to an excitatory subpopulation, cluster, and modulation motif.

#### Linear transformations

We used standardized major axis regression when assessing state-dependent linear transformations, as it adjusts for measurement error on both the x and y axes, which is not accounted for by ordinary least-squares regression [128]. This linear transformation method has been successfully applied in prior work [47, 129]. High and low pupil states were defined as two bins on either side of the median pupil state (62%), whereby the high pupil bin was [72%, 100%] and the low pupil bin was [0%, 52%]. Only sound-responsive neurons with at least 6 stimuli with at least 5 trials per stimulus/state combination across high and low pupil-state bins were used for this analysis. Each neuron’s frequency tuning curve during high trials was regressed onto its respective low-state tuning curve. Slope and y-intercept coefficients were extracted only if they were significantly greater than or less than one or zero, respectively (*α* = 0.05). This was used to determine whether neurons were significantly divisive or multiplicative (based on slope coefficient) and subtractive or additive (based on y-intercept coefficient).

#### Noise and signal correlations

Noise correlations, also known as spike count correlations, were calculated as the Pearson correlation coefficient of the mean-subtracted trial responses between a pair of simultaneously recorded neurons driven by the same stimulus. Trial responses were defined as the mean of raw deconvolved spikes. The mean stimulus response was then subtracted from all trials where that stimulus was presented. These mean-subtracted trial responses were concatenated across all stimuli (15 pure tones) for every neuron in a field of view, yielding a matrix with dimensions [trials x neurons]. The Pearson correlation coefficient was then computed for every neuron pair. Signal correlations, also known as tuning correlations, were computed by taking the frequency tuning curves of a pair of simultaneously recorded neurons and extracting their Pearson correlation coefficient.

All correlation analyses were conducted on sound-responsive neurons and trial activity was defined as the response recorded during a 333 ms evoked response window following stimulus onset. To curtail biased correlations, neurons with a BF of 4 or ≈45.3 kHz were excluded from these analyses. When pupil states were considered, two criteria were applied to include a field of view: (1) neurons must have at least 5 trials at a given stimulus/state combination, and (2) there must be at least 6 stimuli that meet the first criterion.

#### Stimulus decoding

A feed-forward artificial neural network (MATLAB function fitcnet) was trained to classify stimulus identity based on population neural activity at distinct pupil-linked arousal states. The network consisted of an input layer representing population neural activity on each trial, two hidden layers with 16 neurons each, and an output layer with 15 neurons, each corresponding to the predicted probability of one stimulus class. The input to fitcnet was a [trials x neurons] matrix and a one-dimensional vector of stimulus labels (frequency) for each trial. Neural activity for each neuron was defined as its mean trial-evoked response. All decoding was performed separately for each combination of FoV, imaging day, and pupil state. This approach was adopted to maximize the number of neurons used for decoding, as not all neurons within a FoV are consistently observed across all imaging days (e.g., a neurons recorded on Day 1 may not appear on Day 3).

To ensure comparability across conditions, we fixed the number of neurons to 50 and the number of trials per stimulus/state combination to 5 across all decoding runs. When more than 50 neurons and 5 trials per stimulus were available, we performed random resampling within each FoV to create multiple independent decoding sets. Classification accuracy was estimated using 3-fold stratified cross-validation, chosen to accommodate the limited number of trials per class while preserving a balance between training and testing sets. Final performance was quantified by averaging classification accuracy across folds and resampled sets, separately for each pupil state. Decoding was performed independently for each pupil state.

#### Reliability

We used principal component analysis (PCA) in computing population-level reliability to capture network activity across all neurons in a FoV, irrespective of sound responsiveness. To assess tuning curve variance, we first performed PCA on the same [trials x neurons] matrix used as input to the fitcnet network, yielding a [trials x PCs] matrix. We retained only the PCs which cumulatively explained at least 85% of the total variance for our analyses. Then we averaged the trial-level data from this matrix across all trials corresponding to each stimulus, yielding a [stimulus x PCs] matrix, and calculated the variance along the first dimension. To assess trial variance, we calculated the variance along the first dimension of our [trials x PCs] matrix. Reliability for each PC was computed as the ratio of the tuning curve variance to the trial variance, yielding a vector with length of PCs. Finally, we took the mean of this PC vector to get a single reliability value for a given FoV.

Single-neuron reliability was calculated by taking the variance of a neuron’s mean evoked response to each pure tone (tuning curve variance) and dividing it by the variance of that cell’s evoked response at each trial across all stimuli (trial variance) [130]. Mean evoked trial responses were defined as the mean activity during a 333 ms response window following stimulus onset. Single-neuron reliability was conducted on all neurons in a FoV and averaged together, except for **Supplementary Figure 3C**, which was conducted on sound-responsive neurons only and not averaged.

### Statistical Analysis

All statistical analysis was conducted in GraphPad Prism (v10.4.1; GraphPad Software) and R (v4.4.3; R Core Team). Data are reported as mean ± s.e.m. unless otherwise stated. Nonparametric statistical tests were used where data samples did not meet the assumptions of parametric statistical tests. Mixed-effects models were used to assess the effects of fixed factors (e.g., pupil state, excitatory subtype, and inter-somatic distance) on neural activity, with either neuron identity or FoV included as the random effect, depending on the analysis (R packages: lme4, lmerTest, and emmeans). Post-hoc comparisons were conducted using Satterthwaite’s method with Tukey correction for multiple comparisons. Other statistical tests were also corrected for multi-ple comparisons by using Tukey or Dunn’s method, as appropriate. Exact p values were listed if *p <* 10^−100^. Significance on figures denoted as follows: * *p <* 0.05, ** *p <* 0.01, *** *p <* 0.0001.

## Acknowledgements

We thank current and former members of the Williamson Lab for helpful feedback and discussions and assistance with animal care. This work was supported by NIH/NIDCD grants R21DC018327 and R01DC020459, a Hearing Health Foundation Emerging Research Grant, and the Klingenstein-Simons Fellowship in Neuroscience to RSW. Further support was provided by NIH/NIDCD grants T32DC011499 and F31DC021867 to KJK.

## Author Contributions

KJK and RSW conceptualized all experiments. KJK collected and analyzed all data. RFK assisted with software engineering and data analysis. KJK and RSW prepared figures and wrote the manuscript.

---

**Supplementary Figure 1:**
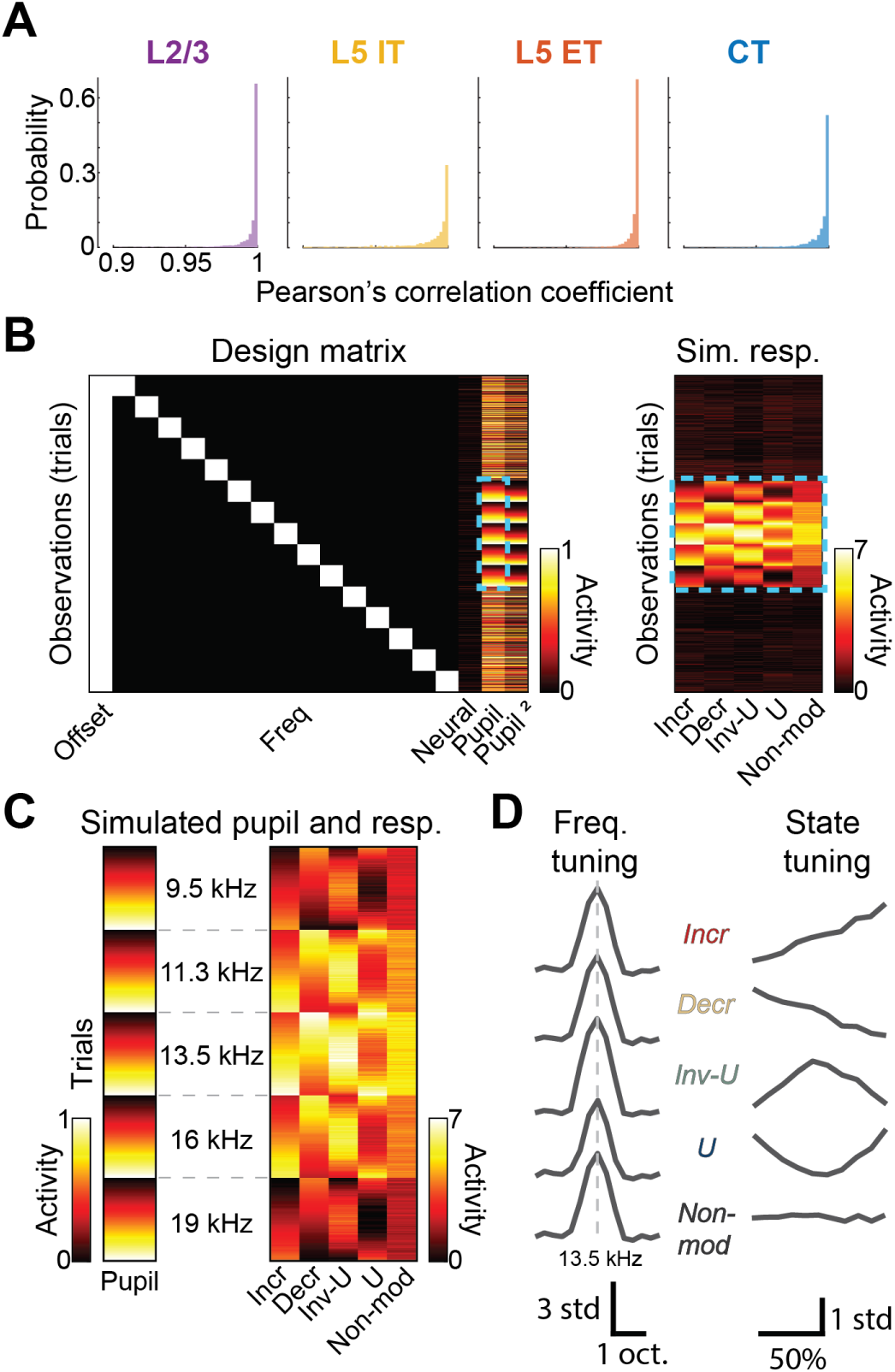
Validation of the multivariate linear regression model. **A**: Probability histogram of the Pearson’s correlation coefficient between a neuron’s frequency tuning curve and its corresponding stimulus coefficients (*β_s_*) from the linear regression model. Distributions significantly differed from a null distribution for each neuron subpopulation (two-sample Kolmogorov–Smirnov test, *p <* 10^−100^). **B**: Design matrix for the linear model (left) that was used for five simulated neurons, each with a different pupil-modulation motif, based on their simulated evoked response vectors (right). **C**: Zoomed in portion of the simulated pupil trace and evoked response vectors shown in the dashed rectangles in (B). **D**: Frequency tuning curves from each of the five simulated neurons (left) and their corresponding pupil-state tuning curves based on responses at best frequency (right). All five simulated frequency tuning curves shared the same best frequency and were nearly identical, but varied slightly due to Gaussian noise.

**Supplementary Figure 2:**
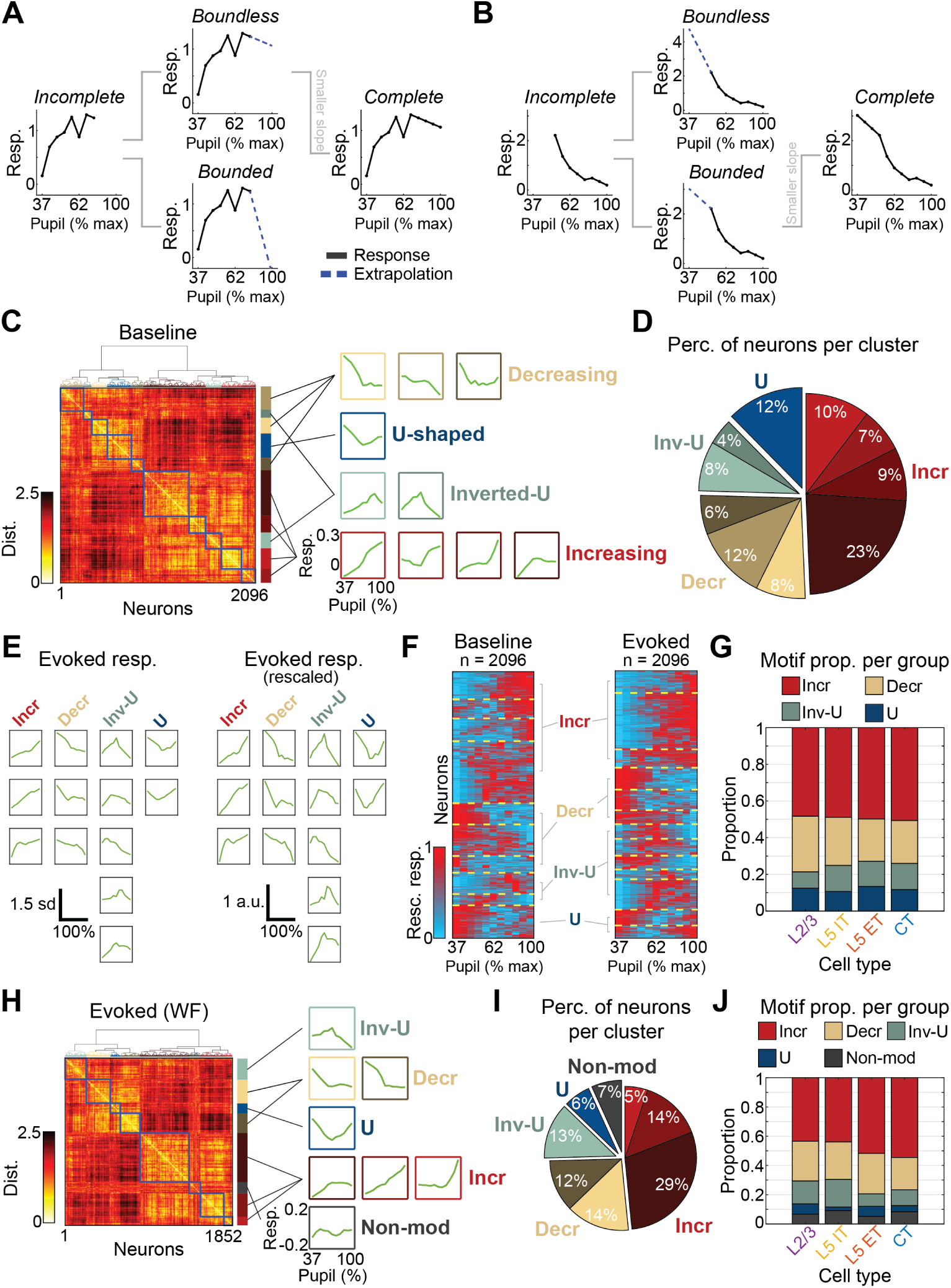
Pupil modulation of baseline and suppressed activity varies by excitatory subtype. **A**: Example of a single neuron with an incomplete BF state tuning curve (left). Comparison of two linear extrapolation methods: boundless and bounded (middle). Boundless extrapolation (dashed blue) yields a shallower slope and was selected to complete the tuning curve (right). **B**: Same as in (A), but for an example where bounded extrapolation is preferred. **C**: Distance matrix from hierarchical clustering of baseline (pre-stimulus) state tuning curves (left). Mean response profiles for each cluster, grouped by modulation motif (right). **D**: Pie chart showing the distribution of modulation motifs across clusters for all excitatory subpopulations. Colors correspond to cluster groupings in (C, right). Modulation motifs were not uniformly represented (*n* = 2096, chi-square goodness-of-fit test, *χ*^2^ = 768.5, *p <* 10^−100^). **E**: Mean z-scored tuning curves by modulation motif (left). Mean rescaled cluster responses organized by modulation motif (right). Each z-scored response is rescaled between 0 and 1. Each motif is shown in the same order across left and right plots. **F**: Color raster plots of mean rescaled state tuning curves across neurons, shown for both baseline (*n* = 2096, left) and evoked (*n* = 2096, right) periods. Yellow dashed lines demarcate clusters, annotated with corresponding modulation motifs. **G**: Proportions of modulation motifs across excitatory subtypes. Motif distributions significantly differed across groups (chi-square test of independence, *χ*^2^ = 20.49, *p* = 0.015). **H**: Same as (C) but clustering was performed on responses to stimuli producing the most suppressed response during the evoked window (WF = worst frequency). **I**: Same as (D) but for WF clustering. Motifs are not uniformly represented (*n* = 1852, chi-square goodness-of-fit test, *χ*^2^ = 1177, *p <* 10^−100^). **J**: Proportions of pupil-state modulation motifs by excitatory subtype. Distribution of motifs significantly differed across groups (chi-square test of independence, *χ*^2^ = 43.44, *p* = 1.9 × 10^−5^).

**Supplementary Figure 3:**
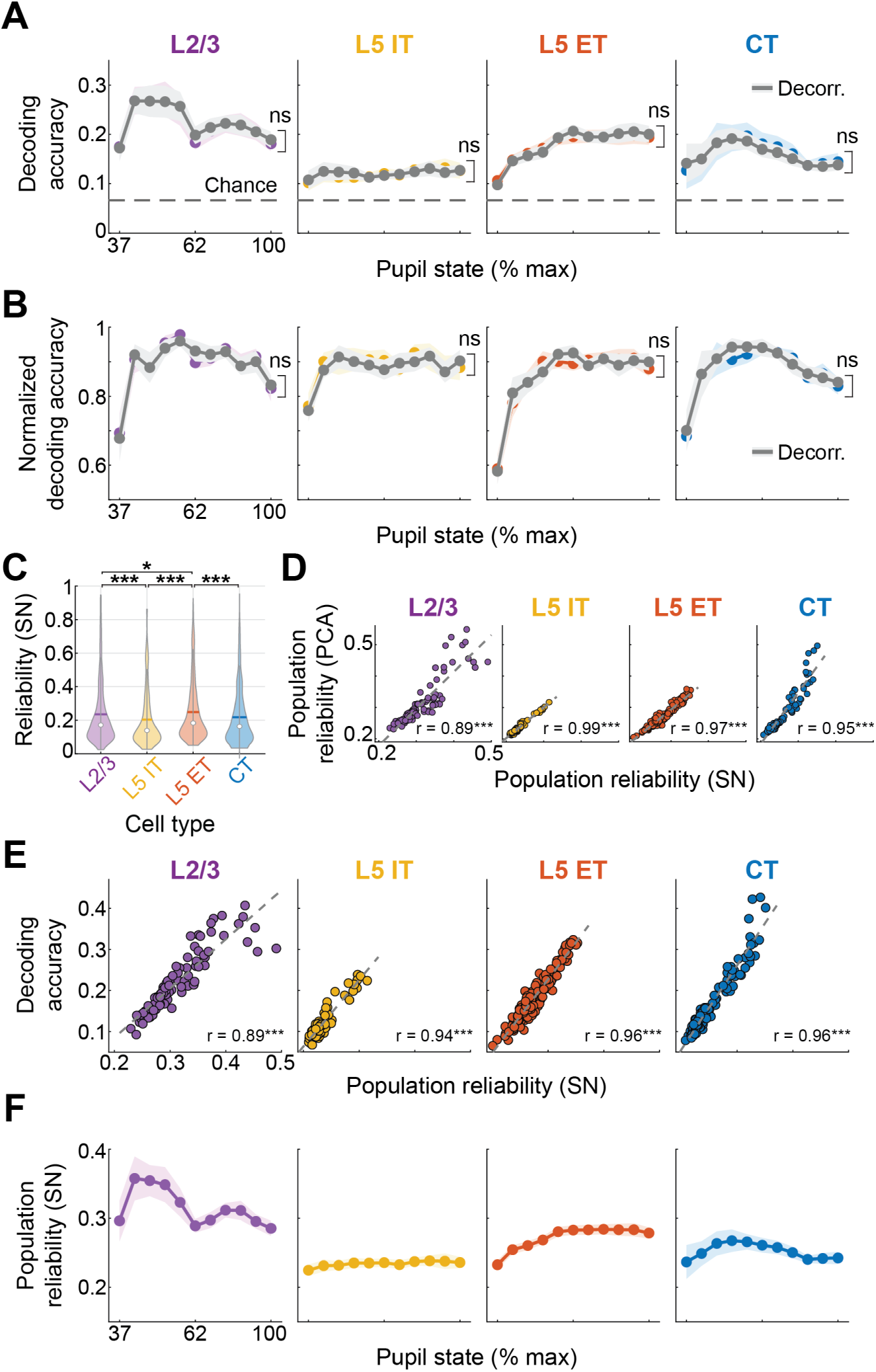
Noise correlations do not affect decoding, while single-neuron reliability predicts decoding performance. **A**: Mean stimulus decoding accuracy across pupil states. Trials within a stimulus class were shuffled to decorrelate neural activity (gray). Decoding performance did not differ between decorrelated and unshuffled datasets (mixed effects model, main effect of shuffling, *p >* 0.99 for all groups). Chance accuracy is 1/15. Shaded regions denote mean ± s.e.m. ns = not significant. **B**: Same as (A), but decoding accuracy was normalized with each FoV to its own maximum value to yield a proportion of max classification performance (value of 1). Decorrelated traces (gray) did not differ from unshuffled data (two-way mixed effects model, *p >* 0.99 for all groups). Shaded regions denote mean ± s.e.m. ns = not significant. **C**: Violin plot of single-neuron reliability for all sound-responsive neurons, irrespective of pupil state. Reliability was not uniform across cell types (*n* = 3336, Kruskal–Wallis test, *χ*^2^ = 42.78, *p* = 2.7 × 10^−9^). White dot: median; bold line: mean. Post-hoc comparisons represented as: * *p <* 0.05, *** *p <* 0.0001. **D**: Scatter plot showing the relationship between PCA-derived and single-neuron population reliability for each FoV at each pupil state. These two reliability metrics were highly correlated for all cell types (L2/3, *r* = 0.89, *p* = 1.7 × 10^−33^; L5 IT, *r* = 0.99, *p* = 7.89 × 10^−83^; L5 ET, *r* = 0.97, *p* = 2.72 × 10^−98^; CT, *r* = 0.95, *p* = 2.01 × 10^−62^). **E**: Same as (D), but comparing mean stimulus decoding accuracy and single-neuron population reliability, which were highly correlated (L2/3, *r* = 0.89, *p* = 4.53 × 10^−34^; L5 IT, *r* = 0.94, *p* = 1 × 10^−51^; L5 ET, *r* = 0.96, *p* = 8.97 × 10^−93^; CT, *r* = 0.96, *p* = 2.24 × 10^−70^). **F**: Mean single-neuron population reliability across pupil states. Reliability was significantly modulated by both pupil state and cell type (two-way mixed-effects model, main effect for pupil and cell type, *F* = 13.59 and 15.79, *p* = 5.17×10^−5^ and *p* = 2.69×10^−8^, respectively; pupil x cell type interaction term, *F* = 2.06, *p* = 0.0012). Shaded regions denote mean ± s.e.m.

